# Disruption of a six-nucleotide miRNA motif improves *PKD1* dosage and ameliorates polycystic kidney disease

**DOI:** 10.1101/2025.10.22.683929

**Authors:** Ronak Lakhia, Chunzi Song, Laurence Biggers, Maggie Zumwalt, Jesus Alvarez, Arvind Somasundaram, Harini Ramalingam, Patricia Cobo-Stark, Vishal Patel

## Abstract

Disrupting microRNA interactions to restore protein expression from haploinsufficient genes offers a promising precision-therapy strategy for monogenic disorders. *PKD1* heterozygosity underlies autosomal dominant polycystic kidney disease (ADPKD), a disorder affecting nearly 12 million people worldwide, where reduced *PKD1* dosage drives progressive cyst formation and kidney failure. We previously identified a 55-bp cis-repressive element in the *PKD1* 3′UTR. Here, we define a six-nucleotide miR-17 seed match within this element that is sufficient to reproduce *PKD1* repression. In vivo base substitution of this motif stabilizes *Pkd1* mRNA and increases polycystin-1 (PC1) protein levels, producing a robust reduction in cyst growth and preservation of kidney function in mouse models. To therapeutically recapitulate this effect, we developed a steric-blocking oligonucleotide that occludes the motif, stabilizes *PKD1* transcript levels, increases PC1 expression, and mitigates cyst-pathogenic events in both murine and patient-derived ADPKD cells. Together, these findings establish a minimal, targetable cis-regulatory motif and provide proof-of-concept for oligonucleotide-mediated *PKD1* derepression, while offering a potentially generalizable strategy to restore other haploinsufficient genes.

## INTRODUCTION

MicroRNAs (miRNAs) are ubiquitous post-transcriptional regulators that fine-tune gene expression by base-pairing to short motifs within mRNA 3′-untranslated regions (3′UTRs)^1, 2^. By coordinately causing relatively modest changes in stability or translation of mRNA networks, miRNAs exert broad effects on cellular physiology and disease phenotypes^3, 4^. However, in certain contexts, a single miRNA interaction can play an outsized regulatory role by suppressing a disease-critical transcript, as occurs in monogenic disorders^5^. Despite this potential, nucleotide-level dissection of individual miRNA motifs, which could enable precise therapeutic targeting, remains rare. This gap is particularly consequential in haploinsufficient diseases, where selectively evading miRNA-mediated repression could restore native gene dosage and confer substantial therapeutic benefit while minimizing off-target effects.

ADPKD provides a compelling example of a haploinsufficient disorder in need of such targeted strategies. ADPKD is most often caused by heterozygous loss-of-function mutations in *PKD1*, the gene encoding polycystin-1 (PC1)^6^. Reduced dosage of *PKD1* initiates cyst formation and drives progressive replacement of functional parenchyma, ultimately leading to kidney failure in many patients. Current disease-modifying therapy is limited to tolvaptan, which slows eGFR decline but has tolerability and safety limitations^7, 8^. Thus, approaches that increase endogenous *PKD1* expression in affected cells are of major therapeutic interest.

The miR-17 family has emerged as a central suppressor of *PKD1* and *PKD2* expression. Genetic and pharmacologic inhibition of miR-17 in preclinical ADPKD models increases polycystin levels and slows cyst growth^5, 9–11^. Notably, the anti–miR-17 oligonucleotide farabursen has demonstrated favorable pharmacodynamic effects, biomarker responses, and trends toward reduced kidney growth in early-phase clinical trials and is now advancing in development for ADPKD^12^. While these studies establish miR-17 as a disease-relevant regulator, they also highlight an alternative strategy—precisely blocking miR-17 interaction on the *PKD1* transcript. This approach would theoretically preserve the broader regulatory functions of the miRNA while selectively restoring the *PKD1* gene.

We previously showed that deletion of a ∼55 bp segment of the *PKD1* 3′UTR that includes a predicted miR-17 site increases *PKD1* translation and attenuates disease phenotypes in ADPKD model systems^5^. However, deletions of this size necessarily remove other cis-elements and potential RNA-binding protein sites, making it difficult to ascribe the observed benefits specifically to the loss of miR-17 binding. Resolving this ambiguity requires nucleotide-level fine mapping to define the minimal motif within the 3′UTR required to mediate miR-17-dependent repression and whether that motif is amenable to targeted therapeutic intervention. Canonical miRNA targeting depends principally on Watson–Crick base pairing between the miRNA seed (nucleotides 2–7) and a complementary stretch in the mRNA^1^. This mechanistic simplicity presents an opportunity for discrete modification or steric masking of those bases to selectively disrupt repression without perturbing broader UTR architecture.

Here, we employ complementary genetic and pharmacologic approaches to address these questions. First, we generated a base-edited allele in which six nucleotides of the *Pkd1* 3′UTR miR-17 motif are altered in situ, preventing canonical pairing while preserving the UTR’s broader architecture. Second, we designed a steric-blocking oligonucleotide complementary to the same minimal motif to test whether transient masking could phenocopy the genetic edit. Using primary renal epithelial cells, engineered cell lines, and mouse models, we evaluated the impact of seed disruption or oligonucleotide masking on *Pkd1*/*PKD1* mRNA stability, PC1 protein abundance, cyst growth, and downstream pathogenic programs.

## RESULTS

### Substitution of six nucleotides in the miR-17 seed motif stabilizes *Pkd1* mRNA

To define the minimal region required for miR-17–*Pkd1* interaction, we targeted six evolutionarily conserved nucleotides within the *Pkd1* mRNA complementary to the miR-17 seed sequence (nucleotides 2–7). Using Cas9/sgRNA-mediated homology-directed repair, we generated a base-substituted allele (*17) with purine-to-purine (A↔G) and pyrimidine-to-pyrimidine (C↔T) substitutions designed to disrupt Watson–Crick base pairing with miR-17 while preserving nucleotide class. This precise modification maintains the overall UTR architecture, allowing us to isolate the effect of this single miRNA interaction site. We introduced this modification into the hypomorphic *Pkd1*^RC3277^ allele, enabling the simultaneous assessment of the cis-regulatory impact on *Pkd1* expression and ADPKD progression ^13^.

We injected *Pkd1*^RC/RC^ zygotes with Cas9 protein, a *Pkd1* 3′UTR-targeting sgRNA, and a single-stranded DNA donor template containing the intended motif sequence changes (5’-CACTTT-3’ → 5’-TGTCCC-3’) within the eight-base-pair miR-17 binding site (Figure 1A). The embryos were implanted in pseudo-pregnant females, and founder mice were genotyped via allele-specific PCR and confirmed via Sanger sequencing (Figures 1B-C). We selected mice carrying one allele with the native miR-17 motif (*Pkd1*^RC^) and the other with the base-edited motif (*Pkd1*^RC*17^), enabling direct allele-to-allele comparisons in the same cells and under a constant genetic background.

**Figure 1:**
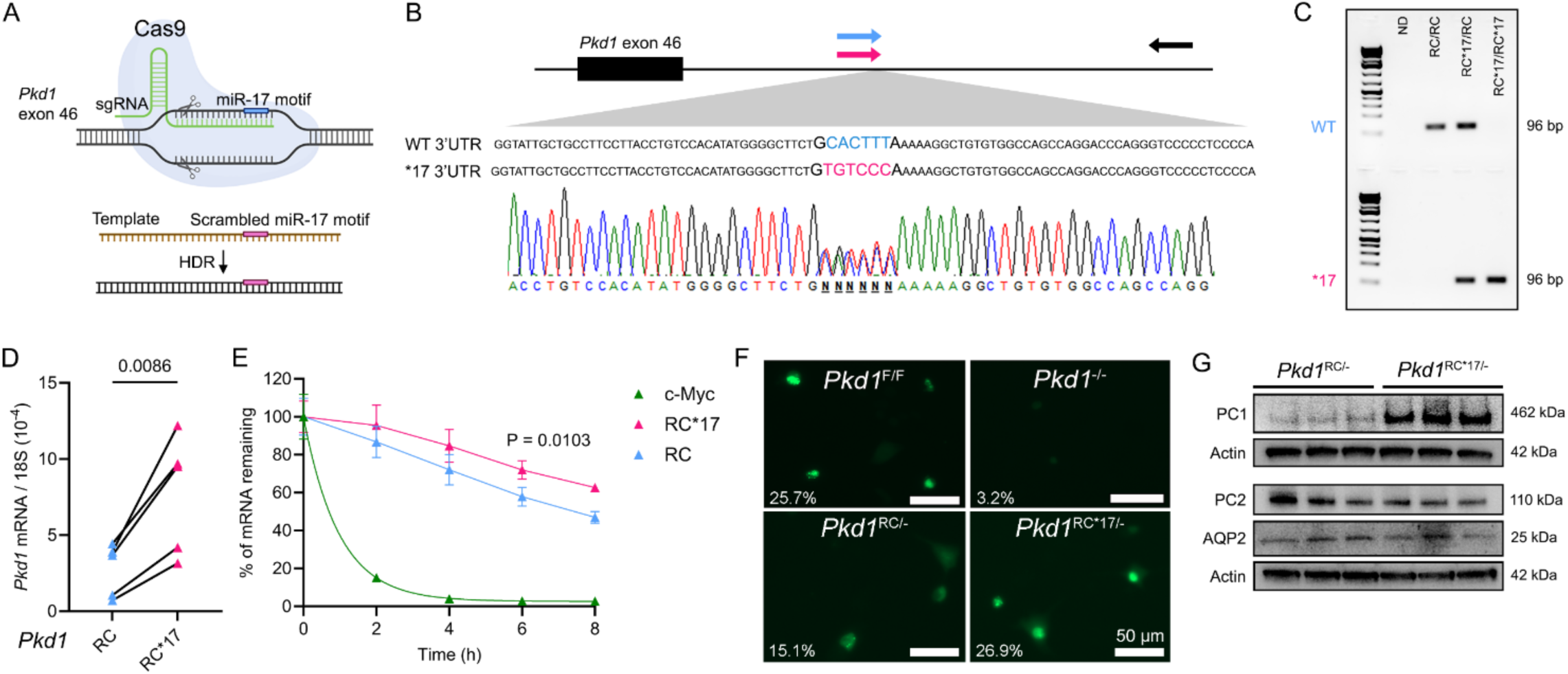
Substitution of six nucleotides within the miR-17 motif stabilizes *Pkd1* mRNA. **A.** Schematic illustrates the CRISPR/Cas9 editing + HDR template DNA approach utilized to precisely switch six nucleotides within the miR-17 motif in the *Pkd1*^RC^ 3’UTR. **B.** Sanger sequencing of tail DNA from *Pkd1*^RC/RC*17^ mouse demonstrates successful 3’UTR editing. The edited allele with disrupted miR-17 motif is denoted in pink, and the unedited allele with intact miR-17 motif is shown in blue. **C.** Allele-specific DNA genotyping primers selectively detected edited (primer positions indicated as pink and black arrows in B) and unedited (primer positions indicated blue and black arrows in B) *Pkd1*^RC^ alleles. **D.** Allele-specific qRT-PCR primers were used to compare *Pkd1*^RC^ and *Pkd1*^RC*17^ transcript abundance within primary kidney epithelial cells derived from *Pkd1*^RC/RC*17^ mice. qRT-PCR demonstrated higher abundance of *Pkd1*^RC*17^ mRNA compared to *Pkd1*^RC^ mRNA. **E.** Primary *Pkd1*^RC/RC*17^ cells were treated with actinomycin and harvested at 2-hour intervals to measure abundance of *Pkd1* mRNA transcripts. qRT-PCR analysis demonstrated slower degradation of *Pkd1*^RC*17^ mRNA compared to *Pkd1*^RC^ mRNA. *c-Myc* mRNA degradation was measured as an internal control. **F.** *Pkd1*^F/F^ or *Pkd1*^−/−^ cells (Top panel) and *Pkd1*^RC/−^ or *Pkd1*^RC*17/−^ cells (lower panel) were transfected with the CRISPR-based *Pkd1* exon-4 mRNA GFP sensor (green). Live-cell images and the quantification of percentage of cells exhibiting the GFP-positive puncta marking *Pkd1* mRNA is shown in the lower left corner. The essential lack of detection of GFP puncta in *Pkd1*^−/−^ cells indicates specificity of *Pkd1* mRNA sensor. A 78% increase in *Pkd1* mRNA detection in *Pkd1*^RC*17/−^ cells compared to *Pkd1*^RC/−^ cells implies increased *Pkd1* mRNA transcript in cells lacking the miR-17 motif. **G.** Western blot demonstrates PC1 expression is increased in *Pkd1*^RC*17/−^ cells compared to *Pkd1*^RC/−^ cells whereas PC2 and AQP2 expression remain unchanged. Error bars indicated SEM. Statistics: Paired t-test (D); nonlinear regression (E); unpaired two-tailed t-test (F).

To test whether the substitution altered endogenous *Pkd1* mRNA stability, we isolated primary kidney epithelial cells from ten-day-old *Pkd1*^RC/RC^, *Pkd1*^RC/RC*17^, and *Pkd1*^RC*17/RC*17^ mice. We designed allele-specific qRT-PCR primers which selectively amplify either the native *Pkd1*^RC^ mRNA or the *Pkd1*^RC*17^ mRNA. The *Pkd1*^RC*17^-specific primer amplified transcripts from *Pkd1*^RC*17/RC*17^ kidney cells but not from *Pkd1*^RC/RC^ kidney cells. Conversely, the *Pkd1*^RC^-specific primer amplified transcripts only from *Pkd1*^RC/RC^ cells but not from *Pkd1*^RC*17/RC*17^ kidney cells (Supplemental Figure 1). Using these primers, we next quantified allele-specific *Pkd1* transcript abundance in the *Pkd1*^RC/RC*17^ mouse kidney epithelial cells. The *Pkd1*^RC*17^ mRNA levels were about threefold higher than *Pkd1*^RC^ mRNA levels (Figure 1D), implying larger contribution to the *Pkd1* mRNA pool from the edited allele compared to the unedited allele. To determine whether the substitution affected transcript stability, we treated *Pkd1*^RC/RC*17^ cells with actinomycin D to halt transcription and harvested RNA every 2 hours for 8 hours. Within the same cells, *Pkd1*^RC*17^ mRNA degraded at a slower rate than the native *Pkd1*^RC^ transcript. After 8 hours, *Pkd1*^RC*17^ mRNA remained significantly more abundant (Figure 1E). We next determined whether the increased mRNA levels due to base substitutions corresponded to higher PC1 in a monoallelic setting. We generated primary and immortalized kidney epithelial cell lines from 14-day-old Ksp^Cre^; *Pkd1*^RC/F^ (*Pkd1*^RC/−^) and Ksp^Cre^; *Pkd1*^RC*17/F^ (*Pkd1*^RC*17/−^) littermate mice. In this model, the floxed *Pkd1* allele is deleted in renal tubules, primarily collecting ducts, generating an ADPKD model with the remaining RC allele that either contains an intact miR-17 motif or the base-edited version. The baseline characterization of these cell lines is shown in Figure 1G and Supplemental Figure 2A-B. First, we used a CRISPR-based live-cell RNA sensing system to detect the *Pkd1*^RC^ transcript. This system employs a catalytically inactive CRISPR-Cas complex (Csm-GFP) with an sgRNA that we designed to specifically recognize and bind exons 2–4 of *Pkd1* mRNA, resulting in GFP fluorescence at the site^14^ (Supplemental Figure 2C). In wildtype cells, we observed punctate GFP signals representing *Pkd1* mRNA, whereas *Pkd1*^−/−^ cells showed no such signals, confirming specificity (Figure 1F). When we employed this RNA sensor system in *Pkd1*^RC/−^ and *Pkd1*^RC*17/−^ cells, we noted a 78% increase in *Pkd1* mRNA signal in *Pkd1*^RC*17/−^ cells. Importantly, immunoblot analysis revealed higher PC1 expression in *Pkd1*^RC*17/−^ compared to *Pkd1*^RC/−^ cells (Figure 1G). Thus, these findings demonstrate that the six-nucleotide substitution within the miR-17 binding site stabilizes the *Pkd1* transcript in cis and leads to higher PC1 protein levels in cultured cells.

### miR-17 motif disruption raises PC1 and ameliorates cyst growth in mice

We next asked whether the *Pkd1*^RC*17^ allele would raise PC1 in vivo and consequently reduce cyst burden in mouse models. To answer this question, we analyzed the impact of miR-17 motif disruption using three independent *Ksp*^Cre^; *Pkd1*^RC/RC*17^ founder mice. In each case, we bred the heterozygous *Ksp*^Cre^; *Pkd1*^RC/RC*17^ founder mice to *Pkd1*^F/F^ mice. This cross produced littermate *Ksp*^Cre^; *Pkd1*^RC/F^ (*Pkd1*^RC/−^) and *Ksp*^Cre^; *Pkd1*^RC*17/F^ (*Pkd1*^RC*17/−^) mice, enabling direct comparison under a similar genetic background. Consistent with our previous studies, the *Pkd1*^RC/−^ progeny from each of the three founder mice developed severe PKD by post-natal day (P)18. In contrast, we noted a marked improvement in cyst burden in their littermate P18 *Pkd1*^RC*17/−^ mice (Figure 2A,E,I). We observed near normalization of kidney-weight-to-body-weight ratio (KW/BW), cyst index, and serum creatinine levels in P18 *Pkd1*^RC*17/−^ compared to *Pkd1*^RC/−^ mice (Figure 2B-C,F-G,J-K, Supplemental Figure 3). Crucially, immunoblot analysis confirmed that P18 *Pkd1*^RC*17/−^ kidneys consistently showed increased PC1 protein abundance compared to *Pkd1*^RC/−^ kidneys across all three founder lines (Figure 2D,H,L).

**Figure 2:**
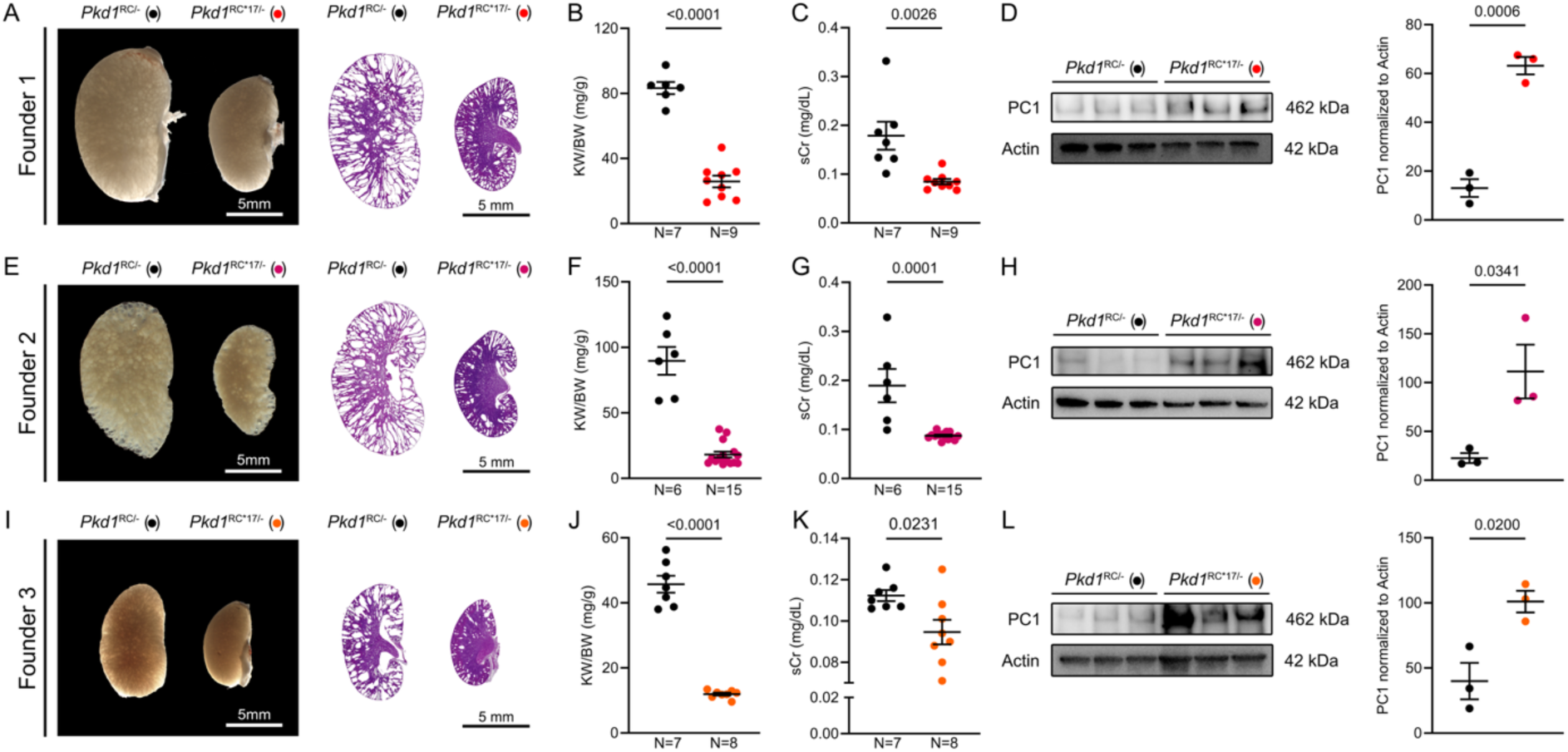
Substitution of six nucleotides within the miR-17 motif ameliorates PKD. Three independent *Pkd1*^RC*17/RC^ founder mouse lines were crossed with Ksp^Cre^; *Pkd1*^F/F^ mice to produce littermate Ksp^Cre^; *Pkd1*^RC*17/F^ (*Pkd1*^RC*17/−^, red circles) and Ksp^Cre^; *Pkd1*^RC/F^ (*Pkd1*^RC/−^, black circles) mice. Representative gross kidney images and H&E-stained kidney sections (**A**), kidney-weight–to–body-weight ratios (KW/BW) (**B**), serum creatinine levels (**C**), and PC1 immunoblot with quantification (**D**) are shown for 18-day-old *Pkd1*^RC*17/−^ and *Pkd1*^RC/−^ progeny derived from founder 1. Equivalent analyses for progeny from founder 2 (**E–H**) and founder 3 (**I–L**) also revealed reduced cystic burden and increased PC1 expression in *Pkd1*^RC*17/−^ mice compared to *Pkd1*^RC/−^ mice. Error bars indicate SEM. Statistical analysis: Unpaired two-tailed t-test. N indicates biological replicates.

To assess durability of the cyst suppression phenotype, we longitudinally followed cohorts of *Pkd1*^RC*17/−^ and *Pkd1*^RC/−^ littermate mice for 3 months. We measured serum creatinine at P45, P60, and P90, performed MRI to measure total kidney volume at P84, and performed histological and molecular analyses at P90. Compared with *Pkd1*^RC/−^ littermates, which showed progressive cyst growth, kidney tissue destruction, and declining renal function, *Pkd1*^RC*17/−^ mice exhibited an attenuated disease course, with preserved renal parenchyma, serum creatinine remaining at near-normal levels, reduced total kidney volume by MRI, and lower KW/BW and cyst index at endpoint (Figure 3A-D, Supplemental Figure 4). Furthermore, we noted that *Pkd1*^RC*17/−^ kidneys have reduced inflammation evidenced by downregulation of MRC1 expression by western blot analysis (Figure 3E) and blunted expression of inflammatory marker transcripts (*Mrc1*, *Ccl2*, *Arg1*, and *Ym1*) by qRT-PCR (Figure 3F). Similarly, we observed a reduction in fibrosis markers, including SMA by western blot analysis (Figure 3G), as well as decreased *Acta2*, *Col1a1*, *Tgfb2,* and *Vimentin* transcripts by qRT-PCR (Figure 3H). Together, these studies confirm the protective effect of a six-nucleotide substitution in the *Pkd1* 3′UTR in delaying cyst progression, and mitigating cyst-associated inflammation and fibrosis.

**Figure 3:**
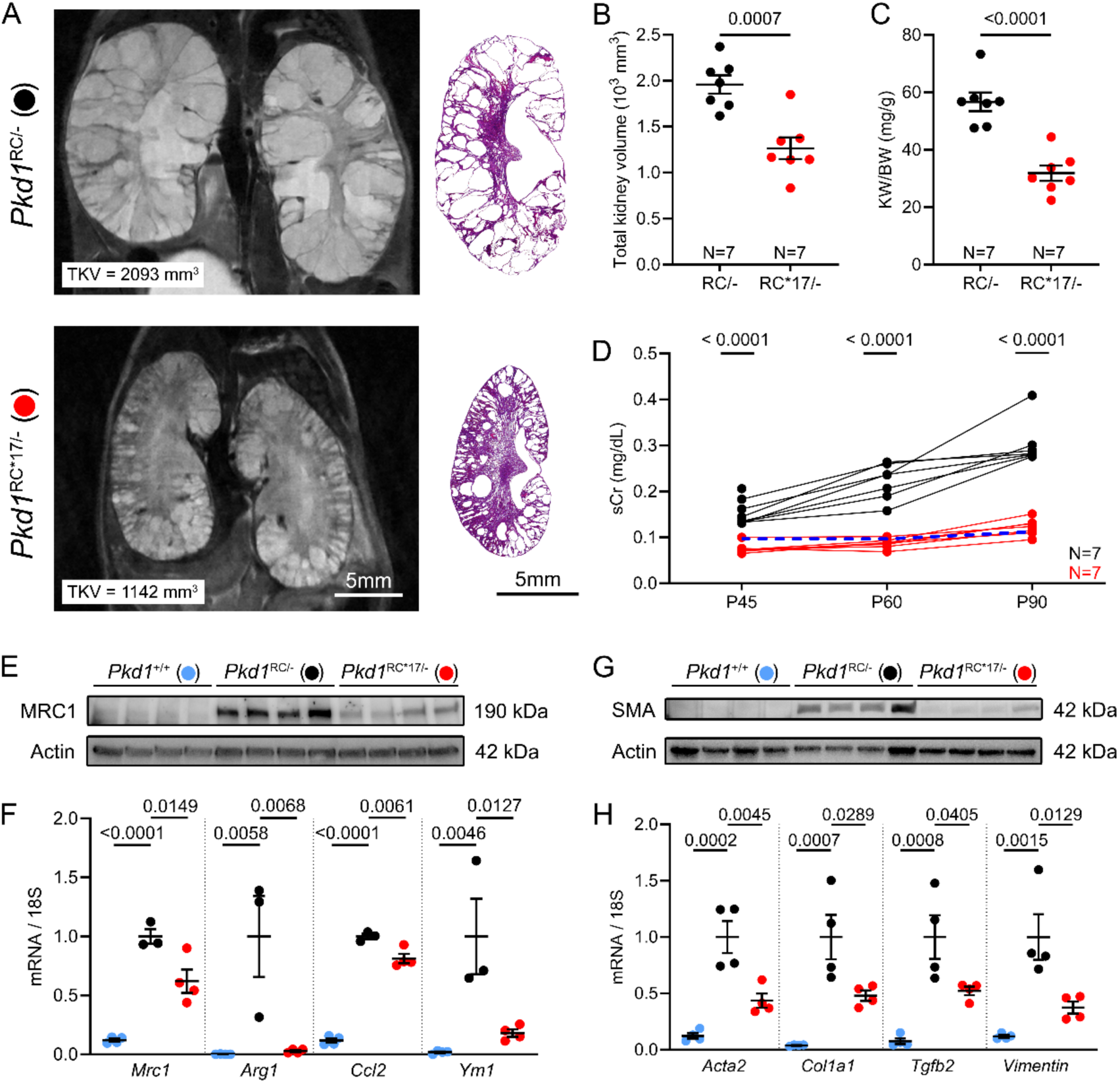
*Pkd1*^RC*17^ substitution results in durable suppression of disease progression. A cohort of *Pkd1*^RC/−^ and *Pkd1*^RC*17/−^ littermate mice derived from founder 1 were aged past postnatal day (P) 18 to examine the durability of disease suppression. MRI at P84 and H&E at P90 (**A**), total kidney volume (**B**), and KW/BW (**C**) demonstrate that cyst growth is significantly retarded even in aged 3-month-old *Pkd1*^RC*17/−^ (red circles) compared to *Pkd1*^RC/−^ mice (black circles). (**D**) Serum creatinine levels at P45, P60, and P90 in *Pkd1*^RC*17/−^ and *Pkd1*^RC/−^ mice are shown. *Pkd1*^RC/−^ mice exhibited a steady increase in serum creatinine with age implying worsening renal function. In contrast, serum creatinine levels remained at or near normal levels (dashed blue line indicates average serum creatinine of wild-type mice; N=4 at each time point) in *Pkd1*^RC*17/−^. Immunoblot (**E** and **G**) and qRT-PCR analysis (**F** and **H**) showing expression of inflammatory markers (**E-F**) MRC1/*Mrc1*, *Arg1*, *Ccl2*, and *Ym1* and fibrosis markers (**G-H**) SMA/*Acta2*, *Col1a1*, *Tgfb2*, and *Vimentin* in kidneys of *Pkd1*^+/+^, *Pkd1*^RC/−^, and *Pkd1*^RC*17/−^ are shown (N=4 for each group). Error bars indicate SEM. Statistical analysis: Unpaired, two-tailed t-test (B-D); ANOVA with Tukey’s (F-H).

### PC1 restoration disrupts the c-Myc–miR-17–mitochondrial axis in ADPKD

Our miR-17 motif–edited *Pkd1* allele provides a powerful system to address a central unresolved question: what are the downstream molecular consequences of restoring endogenous PC1? Mitochondrial dysfunction and hyperproliferation, orchestrated in part by the c-Myc–miR-17 axis, are major drivers of ADPKD, yet it remains unclear the extent to which PC1 restoration can directly reprogram these pathogenic processes^15–17^. Because this allele uncouples PC1 from miR-17 regulation, it offers a unique opportunity to isolate the effects of PC1 restoration on mitochondrial function and c-Myc signaling.

To obtain converging lines of evidence, we simultaneously assessed global proteomic changes, cellular proliferation, and mitochondrial function in kidneys from P18 *Pkd1*^RC*17/−^ mice and *Pkd1*^RC/−^ mice. Shotgun label-free quantitative proteomics revealed widespread changes in protein abundance, with 733 proteins upregulated and 1624 proteins downregulated in *Pkd1*^RC/−^ relative to age-matched non-cystic *Pkd1*^+/+^ control kidneys (Supplemental Figure 5 and Supplemental Table 1). Nearly 50% of dysregulated proteins in *Pkd1*^RC/−^ kidneys showed improved (or normalized) expression *Pkd1*^RC*17/−^ kidneys, implying that along with producing a marked improvement in disease burden, PC1 restoration also positively affects the downstream proteo-molecular footprint (Figure 4A). Pathway enrichment analysis highlighted mitochondrial metabolism, cell proliferation, and RNA translation and processing as the major processes that were positively affected by miR-17 motif disruption (Figure 4B). Consistent with these findings, immunoblotting using a panel of antibodies demonstrated marked improvement in the expression of mitochondrial electron transport chain complexes in *Pkd1*^RC*17/−^ kidneys compared to *Pkd1*^RC/−^ kidneys (Figure 4C). qRT-PCR analysis further revealed improved expression of *Ppargc1a*, a key regulator of mitochondrial biogenesis, in *Pkd1*^RC*17/−^ compared to *Pkd1*^RC/−^ kidneys, supporting enhanced mitochondrial function (Supplemental Figure 6A). In parallel, we noted reduced expression of p-mTOR and pCREB1 (Figure 4C), implying that global proteomic changes were associated with attenuation of cyst-pathogenic signaling. Supporting this conclusion, whole-section immunofluorescence quantification of phospho-histone H3-positive cells revealed a 19% reduction in cellular proliferation in *Pkd1*^RC*17/−^ kidneys compared to *Pkd1*^RC/−^ kidneys (Figure 4D-E). Collectively, these complementary analyses indicate that PC1 restoration simultaneously enhances mitochondrial function and restrains hyperproliferative, cystogenic programs.

**Figure 4:**
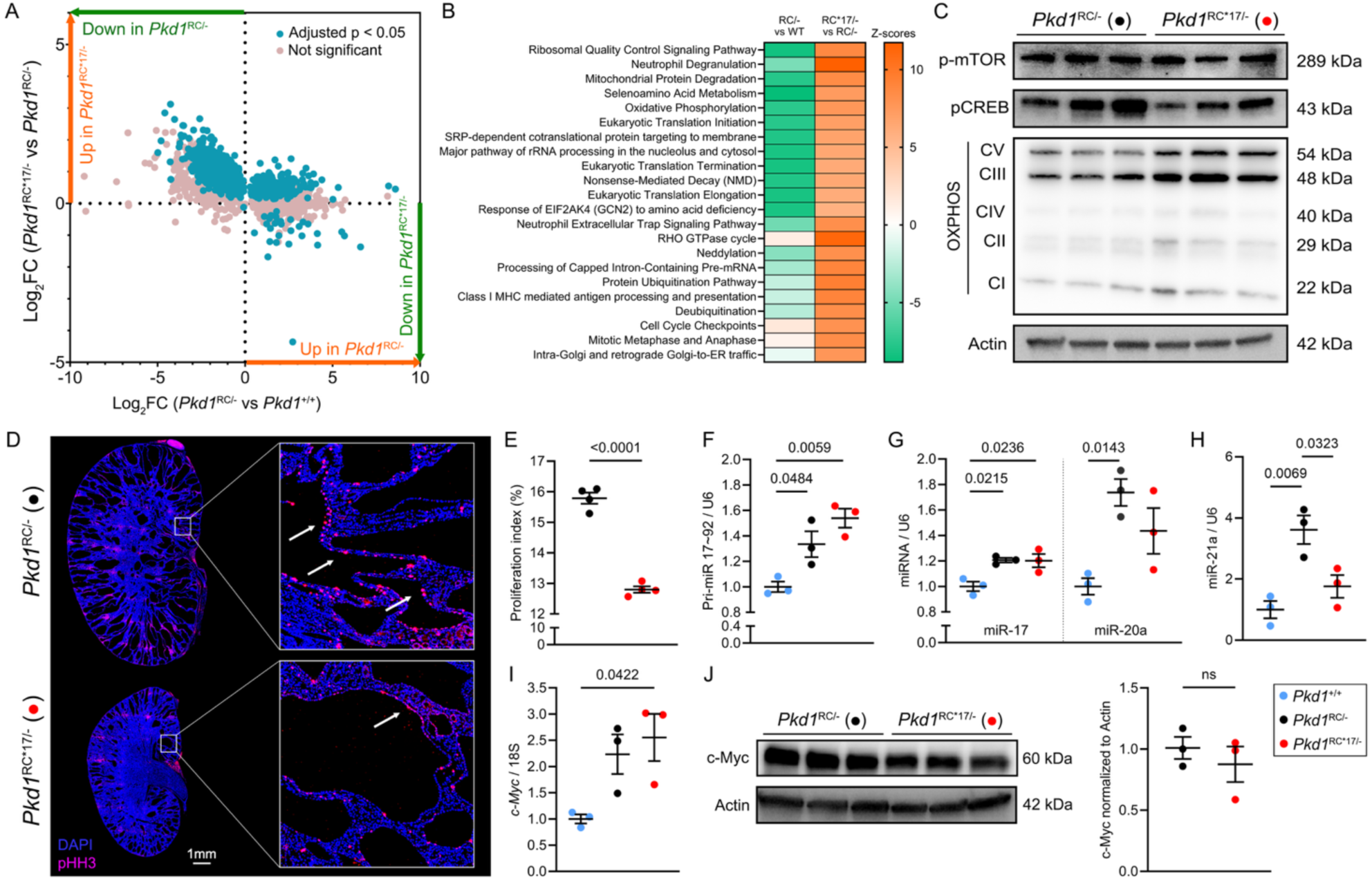
*Pkd1*^RC*17/−^ allele disrupts the c-Myc–miR-17–mitochondrial axis in cystic kidneys. Shotgun label-free quantitative proteomics was performed to assess global proteomic profiles of kidneys from 18-day-old *Pkd1*^+/+^, *Pkd1*^RC/−^, and *Pkd1*^RC*17/−^ mice (n=3, each group). **A.** Comparative differential expression analysis depicts a global trend where a substantial portion of dysregulated protein expression (adjusted p <0.05) in *Pkd1*^RC/−^ kidneys compared to *Pkd1*^+/+^ kidneys was improved in *Pkd1*^RC*17/−^ kidneys. **B**. Top pathways (based on *z*-score ≥ 6 in at least one comparison group) that underlie the proteomic changes observed in *Pkd1*^RC/−^ compared to *Pkd1*^+/+^ kidneys, and *Pkd1*^RC*17/−^ compared to *Pkd1*^RC/−^ kidneys. The ingenuity pathway analysis software was used for the predictions. Expression of metabolic, proliferative, and RNA processing pathways were improved in *Pkd1*^RC*17/−^ compared to *Pkd1*^RC/−^ kidneys. **C.** Immunoblot analysis showing reduced pMTOR and pCREB expression, and increased expression of oxidative phosphorylation complex (OXPHOS) components in *Pkd1*^RC*17/−^ compared to *Pkd1*^RC/−^ kidneys (n=3). **D-E**. Phospho-histone H3 (pHH3, pink) immunofluorescence staining (**D**) and subsequent quantification (**E**) demonstrated reduced proliferation in *Pkd1*^RC*17/−^ compared to *Pkd1*^RC/−^ kidneys (n=4 each). The slides were counterstained with DAPI (blue). **F-I.** qRT-PCR analysis showing expression of primary miR-17 transcript (**F**), mature miR-17 and miR-20a (**G**), mature miR-21 (**H**), and *cMyc* (**I**) in kidneys of 18-day-old *Pkd1*^+/+^, *Pkd1*^RC/−^, and *Pkd1*^RC*17/−^ mice (n=3). The c-Myc/miR-17 axis remained elevated in *Pkd1*^RC*17/−^ kidneys, whereas expression of miR-21, a cyst-pathogenic miRNA, was improved. **J.** Immunoblot analysis showing equivalent expression of cMyc protein in kidneys of 18-day-old *Pkd1*^RC/−^ and *Pkd1*^RC*17/−^ mice (n=3). Actin serves as the loading control; Error bars indicate SEM. Statistics: ANOVA with Tukey’s (A, F-I), and Unpaired, two-tailed t-test (E, J)

Finally, we asked whether PC1 restoration influenced the c-Myc–miR-17 axis itself. c-Myc is broadly activated in ADPKD and was one of the first oncogenes implicated in the disease, driving both cellular proliferation and the expression of miR-17 family microRNAs^11, 15^. As expected, pri-miR-17 was transactivated at 33% higher levels in *Pkd1*^RC/−^ kidneys compared with non-cystic controls (Figure 4F). Remarkably, pri-miR-17 levels remained unchanged in *Pkd1*^RC*17/−^ kidneys, and mature miR-17 and miR-20a remained similarly elevated (Figure 4F-G). Correspondingly, *c-Myc* mRNA and protein levels were unchanged in *Pkd1*^RC*17/−^ kidneys compared with *Pkd1*^RC/−^ kidneys (Figure 4I-J). In contrast, the expression of miR-21, another cystogenic miRNA upregulated *Pkd1*^RC/−^ kidneys, was reduced in *Pkd1*^RC*17/−^ kidneys (Figure 4H), suggesting that PC1 restoration selectively modulates certain cystogenic microRNAs while leaving the c-Myc–miR-17 axis largely intact^18^. These findings indicate that the beneficial effects of PC1 restoration on cyst growth, mitochondrial function, and cellular proliferation occur independently of c-Myc and miR-17, highlighting a direct role of PC1 in rewiring downstream pathogenic programs.

### Design and validation of *PKD1*-targeting antisense oligonucleotides

Having established that precise base editing of the miR-17 seed in *Pkd1* stabilizes its mRNA and elevates PC1, we next asked whether a transient, pharmacologic approach could recapitulate these effects. To test this, we employed a multifaceted strategy to determine the optimal anti-sense oligonucleotide (ASO) sequence capable of masking the miR-17 binding motif (Supplementary Figure 7, Supplemental Table 2,3). Our design process integrated four key criteria: 1) maximization of hybridization, by analyzing the predicted secondary RNA structure of the *Pkd1*/*PKD1* 3’UTR to target the motif within an accessible open loop; 2) minimization of off-targets, achieved by a genome-wide analysis that identified 15-16 bp length as optimal for specificity in both mouse and human; 3) inter-species conservation, to ensure the oligo sequence was effective in both mice and humans; and 4) sequence stability, by screening candidates to select for those with the least potential for self-dimerization and hairpin formation. This multi-parameter optimization led us to two lead candidates: 16 bp mouse *Pkd1*-oligo (P1) and 15 bp human *PKD1*-oligo (P2). The P2 oligo is fully complementary to the human *PKD1* 3’-UTR with one base mis-match to the corresponding mouse *Pkd1* 3’UTR. The complement of the miR-17 motif resides proximal to 5’ end of the P1 and P2 oligos. The sequences are shown in Supplementary Figure 7. Importantly, these oligos are fully modified DNA oligonucleotides designed to promote steric hindrance of miRNA binding rather than mRNA cleavage via RNase H.

First, we confirmed target engagement by exploiting the property that RNA-oligo duplexes physically obstruct reverse transcriptase during cDNA synthesis^19^. We transfected mouse collecting duct cells with 40 nM of either P1-oligo or a scrambled control oligonucleotide (Ctl-oligo) and measured *Pkd1* mRNA levels via qRT-PCR 48 hours post-transfection. The P1-oligo prevented amplification of the 3′UTR region it targets, while adjacent or unrelated regions remained detectable, indicating specific binding to the intended motif (Supplemental Figure 8). This hindrance was not observed in cells treated with Ctl-oligo. To demonstrate functional engagement, we co-transfected a luciferase reporter fused to the *Pkd1* 3′UTR with either a control or miR-17 mimic in mouse collecting duct cells. As expected, miR-17 suppressed luciferase activity. Co-transfection with P1-oligo mitigated this suppression, restoring reporter expression, whereas Ctl-oligo had no effect (Supplemental Figure 9A). Similar results were obtained in *Pkd1*^RC/−^ cells, confirming that the oligo’s activity is consistent across different cellular contexts (Supplemental Figure 9B). Collectively, these experiments demonstrate that P1-oligo can specifically inhibit miR-17–*Pkd1* interactions at the 3′UTR.

Next, we employed multiple complementary methods to assess whether P1 oligo binding stabilizes the endogenous *Pkd1* mRNA. First, we treated normal kidney collecting duct cells with 40 nM of either Ctl-oligo or P1 oligo for 48 hours, followed by treatment with actinomycin D to inhibit new RNA synthesis. The actinomycin D chase assay revealed that cells treated with P1 oligo retained approximately 30% more *Pkd1* mRNA after 6 hours compared to Ctl-oligo-treated cells, indicating enhanced transcript stabilization (Supplemental Figure 10A-B). As an orthogonal validation, we used our *Pkd1* RNA sensing system^14^. We measured *Pkd1* mRNA abundance in live cells treated with 40 nM of P1 oligo or Ctl-oligo. After 48 hours, P1-treated cells exhibited a significant increase (approximately 75%) in *Pkd1* mRNA compared to Ctl-oligo-treated cells (Supplemental Figure 10C). Finally, immunoblot analysis revealed that the stabilized *Pkd1* mRNA translates into ∼40% increased PC1 protein with P1 oligo compared Ctl-oligo treatment (Supplemental Figure 10D). Together, these findings demonstrate that steric hindrance of the miR-17 motif via oligonucleotide binding stabilizes endogenous *Pkd1* mRNA, leading to enhanced translation and increased PC1 protein expression.

### Pkd1 oligo restores PC1 expression and reverses cyst-pathogenic events in ADPKD models

While studies in normal cells established that the P1 oligo can stabilize wildtype *Pkd1* transcripts, the critical question was whether the same strategy could rescue PC1 expression and cyst-pathogenic events in monoallelic, disease-relevant, mouse and human ADPKD cells. To address this, we first turned to *Pkd1*^RC/−^ cells. Just as in wildtype cells, the P1 oligo bound the 3′UTR, blocked reverse transcription across the miR-17 motif, and stabilized the *Pkd1*^RC^ mRNA transcripts following actinomycin D treatment (Figure 5A-C). Moreover, in situ transcript quantification in live cells using the Csm-GFP system revealed that *Pkd1* mRNA was reduced by ∼50% in *Pkd1*^RC/−^ compared to *Pkd1*^RC/+^ cells. Treating *Pkd1*^RC/−^ cells with P1 oligo reversed this reduction whereas control oligo had no effect (Figure 5D). Importantly, immunoblot analysis also revealed restored PC1 protein levels in *Pkd1*^RC/−^ cells following P1 oligo treatment compared Ctl-Oligo treatment (Figure 5E). We next asked whether these molecular effects could mitigate pathogenic features of PKD. Indeed, P1-oligo reduced proliferation of *Pkd1*^RC/−^ cells in 2D culture and decreased cyst size in 3D Matrigel assays (Figure 5F). At the signaling level, P1 oligo treatment suppressed the cyst-promoting effector pCREB, and at the metabolic level, it restored mitochondrial membrane potential (Figure 5G). Thus, stabilization of the *Pkd1*^RC^ allele not only restored PC1 expression but also reversed multiple cellular hallmarks of PKD.

**Figure 5:**
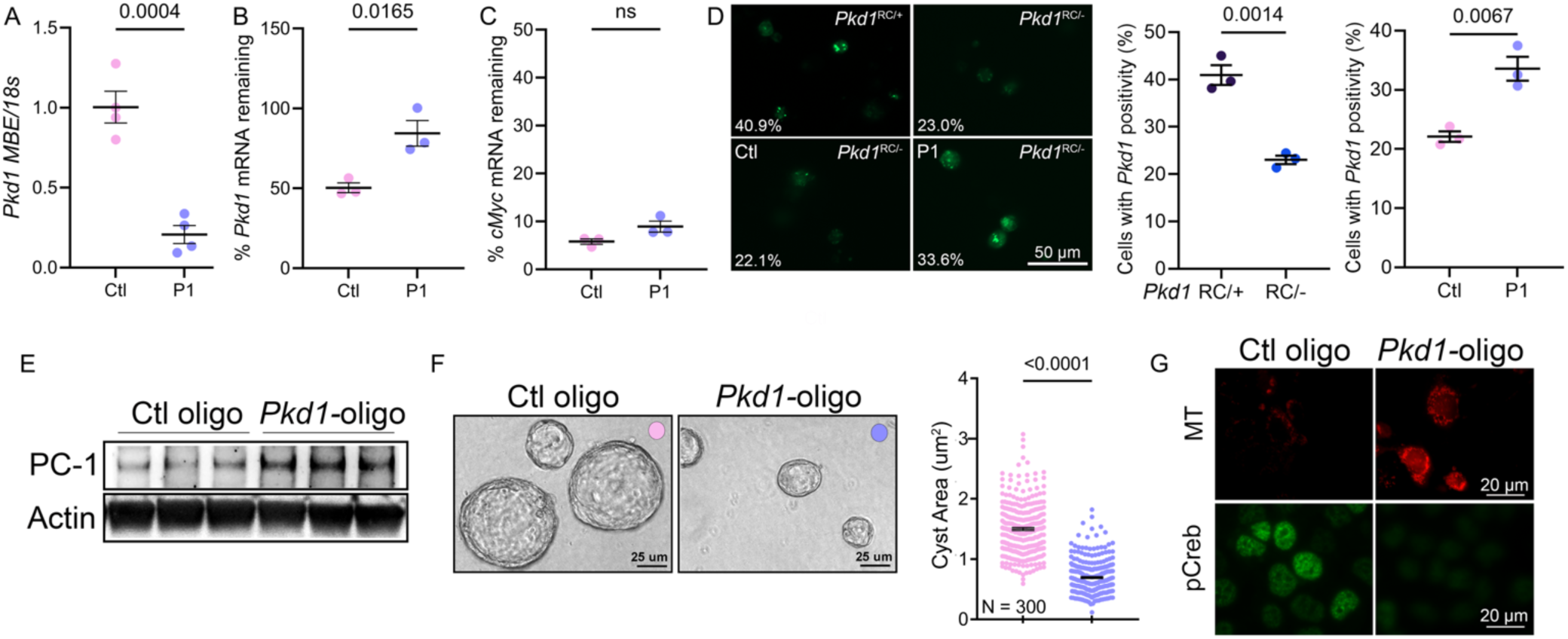
Pkd1-oligonucleotide increases PC1 expression and improves cyst parameters in mouse cellular PKD model. *Pkd1*^RC/−^ cells were treated with 40 nM control oligonucleotide (Ctl oligo) or Pkd1-oligonucleotide (P1) and analyzed 48 hours later. **A.** qRT-PCR analysis was performed using a primer pair that amplifies the *Pkd1 3*’UTR. P1 prevented amplification of the *Pkd1* 3′UTR region it targets, while control oligo had no effect, indicating specific binding of P1 to the intended motif. **B-C.** *Pkd1*^RC/−^ cells were treated with 40 nM Ctl or P1 for 48 hours and then co-treated with actinomycin for four hours to halt *de-novo* transcription. qRT-PCR analysis using a primer pair that amplifies *Pkd1* exon-4 showed that 68% more *Pkd1* mRNA remained in P1-treated compared to Ctl-oligo-treated *Pkd1*^RC/−^ cells whereas *cMyc* transcript degraded equivalently. **D.** (Top panel) *Pkd1*^RC/+^ or *Pkd1*^RC/−^ cells were transfected with the CRISPR-based *Pkd1* exon-4 mRNA GFP sensor (green). (Lower panel) *Pkd1*^RC/−^ cells treated with 40 nM Ctl or P1 oligo and co-transfected with the CRISPR-based sensor. Live cell imaging and quantification of percentage of cells exhibiting the GFP signal is shown. *Pkd1* mRNA detection was reduced by 44% in *Pkd1*^RC/−^ cells compared to *Pkd1*^RC/+^ cells. Treatment with P1 oligo improved *Pkd1* mRNA detection by 50% compared to Ctl-oligo treatment. **E.** Immunoblot showing increased PC1 expression in *Pkd1*^RC/−^ cells 48-hr after P1 treatment compared to Ctl treatment. Actin serves as the loading control. **F.** *Pkd1*^RC/−^ cells were transfected with 40 nM Ctl or P1 for 48 hours and were then cultured in 3D Matrigel for five days to facilitate cyst formation. Quantification showing that cyst size was reduced in P1-oligo compared to Ctl-oligo-treated *Pkd1*^RC/−^ cells. **G.** Immunofluorescence images showing live-cell MitoTracker labeling (MT, red) or pCreb (green) expression in P1-treated compared to Ctl oligo-treated *Pkd1*^RC/−^ cells. P1 treatment increased Mitotracker signal and reduced pCreb expression, implying improved mitochondrial function and reduced cAMP signaling. N=3 biological replicates for each condition. Error bars indicate SEM. Statistical analysis: Unpaired, two-tailed, t-test (A-D, F). N=4 biological replicates (A). N=3 biological replicates (B-D).

Finally, we extended these studies to human disease models, and interchangeably tested both P1 and P2 oligos. We examined independent ADPKD cell lines derived from cystic kidney epithelia isolated from nephrectomy specimens of three individuals with ADPKD, and each carrying a distinct pathogenic monoallelic *PKD1* mutation^5^. Immunoblot analysis revealed that both P1 and P2 increased PC1 protein expression across all three patient-derived ADPKD cell lines (*PKD1* Q2556X, *PKD1* Q2641*, and *PKD1* pR3926Afs*34) as well as in the *Pkd1*^RC/–^ mouse line (Figure 6A). qRT-PCR using primers that amplify the 3′UTR miR-17 motif demonstrated that P2 selectively interfered with amplification of the human 3′UTR region, while P1 selectively inhibited the mouse 3′UTR (Figure 6B). Each oligonucleotide showed partial cross-reactivity in the non-cognate species, whereas the control oligo had no effect (Figure 6B). Moreover, in a 3D Matrigel cystogenesis assay, treatment with both P1 and P2 oligo reduced cyst size in two human ADPKD cell lines (Figure 6C). These consistent results across mouse and human systems establish that the P1 and P2 oligo can derepress non-inactivated *PKD1* alleles, restore PC1 expression, and revert disease-associated phenotypes in cellular models. Collectively, they provide a strong rationale for advancing this approach as a therapeutic strategy in ADPKD.

**Figure 6:**
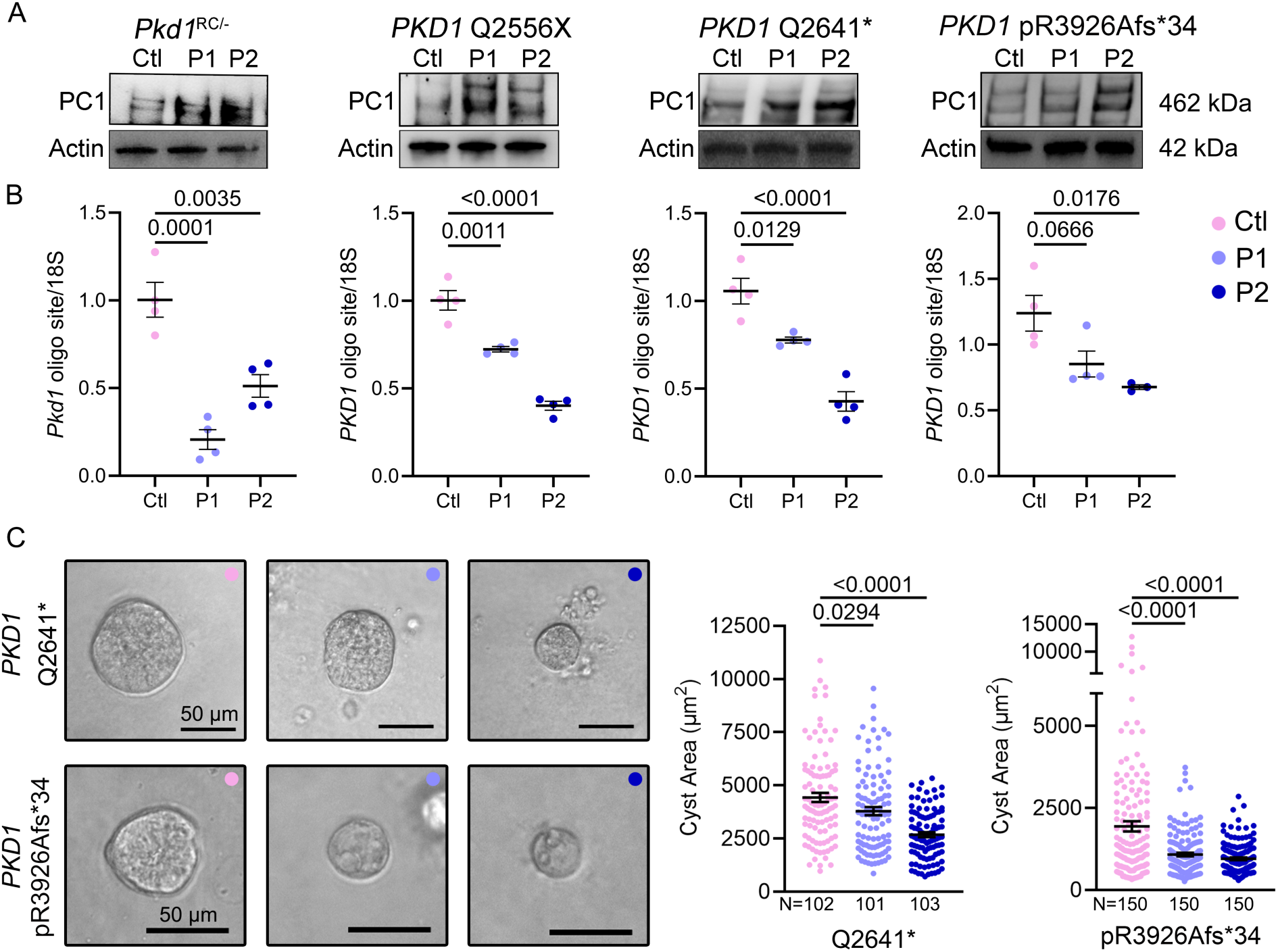
PKD1-oligonucleotides raise PC1 protein levels in patient-derived ADPKD cyst cells. Three independent patient-derived ADPKD cyst cell lines (*PKD1* Q2556X, *PKD1* Q2641*, and *PKD1* pR3926Afs*34) were treated with 40 nM control oligonucleotide (Ctl), the mouse-specific *Pkd1*-targeting oligo (P1), or the human-specific PKD1-targeting oligo (P2). The P1 and P2 oligos were also tested in the mouse *Pkd1*^RC/−^ cell line. **A.** Immunoblot analysis revealed that both P1 and P2 increased PC1 protein expression across all three patient-derived ADPKD cell lines, as well as in the *Pkd1*^RC/–^ mouse line. **B.** qRT-PCR using primers that amplify the *PKD1* 3′UTR miR-17 motif demonstrated that P2 selectively interfered with amplification of the human 3′UTR region, while P1 selectively inhibited the mouse 3′UTR. Each oligonucleotide showed partial cross-reactivity in the non-cognate species, whereas the control oligo had no effect. **C.** Human ADPKD cell lines were treated with 40 nM ctl, P1, or P2 oligos for 48-hours and then cultured in 3D Matrigel. Quantification revealed that treatment with either P1 or P2 oligo reduced 3D cyst size compared to Ctl oligo. Error bars indicate SEM. Statistical analysis: ANOVA with Tukey’s (B-C). N=3 biological replicates for each cell line (B). N=101-150 cysts were measured for each condition (C).

## DISCUSSION

In this study, we define both a mechanistic and therapeutic framework for selectively derepressing *PKD1* by targeting a discrete microRNA-binding motif in its 3′UTR. Through systematic motif mapping, we identify a six-nucleotide stretch that is essential for miR-17– mediated repression of *PKD1*. Although miRNA biology has been studied extensively, fine-mapping of functional cis-elements at nucleotide resolution in mammalian disease genes is rare. By achieving this level of mechanistic granularity in a clinically relevant context, our work establishes a paradigm for pinpointing minimal regulatory sites that can be exploited for therapeutic intervention. Disruption of this site—either genetically, through precise base editing, or transiently, with steric-blocking oligonucleotides—stabilizes *Pkd1*/*PKD1* transcripts, elevates PC1 protein, and attenuates cyst growth in mouse and human models. Importantly, our approach narrows the therapeutic target compared with prior studies that removed broader 3′UTR regions potentially containing multiple cis-elements^5^. By isolating a minimal motif sufficient for oligonucleotide masking, we both refine the mechanistic understanding of PKD1 regulation and provide a tractable strategy for restoring haploinsufficient genes.

Restoring PC1 expression also gates downstream disease pathways. Although c-Myc and miR-17 remain elevated, mitochondrial function and proliferation normalize once PC1 levels are restored. This indicates that it is the loss of PC1 that opens the gate to pathogenic outputs, whereas re-establishing PC1 closes it and prevents downstream hyperproliferation and metabolic dysfunction despite persistent upstream oncogenic signals. In this way, PC1 functions as a central checkpoint that governs proliferative and metabolic balance, and its restoration is sufficient to re-establish cellular homeostasis.

Beyond PKD1, our findings illustrate a broader therapeutic principle. Precise targeting of a minimal cis-regulatory miRNA motif can selectively derepress a disease-critical transcript without globally inhibiting its cognate miRNA. Compared with anti–miR-17 strategies that silence the miRNA network more broadly, motif masking preserves the wider physiological functions of miR-17 while still achieving therapeutic benefit. Moreover, this approach is mutation-agnostic. By stabilizing any residual *PKD1* transcript, it could restore PC1 dosage across diverse allelic backgrounds, bypassing the challenge of correcting thousands of distinct *PKD1* mutations in the patient population. Derepression of *PKD1* through a single regulatory site therefore offers a unifying therapeutic entry point for ADPKD.

The robustness of our findings is supported by the breadth of models tested. We engineered three independent founder mouse lines with a targeted *Pkd1* 3′UTR mutation, each demonstrating PC1 stabilization and cyst suppression. In parallel, steric-blocking oligonucleotides were validated in a normal mouse cell line and in four distinct ADPKD cell models (one mouse and three human), consistently restoring *PKD1* expression and attenuating cystic growth. This convergence across species, alleles, and experimental systems underscores the reproducibility and generalizability of the mechanism.

Several limitations warrant consideration. First, although we demonstrate strong effects on transcript stability, protein restoration, and cyst growth in vitro, in vivo testing of the Pkd1-targeting oligonucleotide remains to be performed. Translation toward therapy will require optimization of oligonucleotide chemistry, dosing, and kidney-targeted delivery to ensure stability, bio-distribution, and sustained engagement. However, this limitation is mitigated by precedence involving our work on anti–miR-17 oligonucleotides RGLS4326 and RGLS8429^10^. These oligos achieve efficient renal delivery and robust target engagement in vivo, supporting the feasibility of applying steric-blocking oligonucleotides to the kidney. Second, this strategy requires at least one functional *PKD1* allele. Many cysts, particularly larger, late-stage ones, have undergone somatic loss of the second allele and would therefore remain refractory to *PKD1*-restoring interventions^20, 21^. However, a recent study showed that up to 50% of cysts in end-stage ADPKD specimens retain residual *PKD1* expression, suggesting that a substantial fraction of cysts could still be responsive to *PKD1*-restoring interventions^22^.

In summary, our work delivers three key insights. First, nucleotide-level motif mapping in a mammalian disease context can reveal minimal cis-regulatory sites whose masking is sufficient to restore haploinsufficient gene expression. This remains a rare but powerful strategy for both mechanistic dissection and therapeutic intervention. Second, PC1 dosage gates downstream pathogenic programs. Low PC1 opens the gate to mitochondrial dysfunction and hyperproliferation, whereas restored PC1 closes it and normalizes cell state despite persistent oncogenic signals. Third, steric-blocking oligonucleotides targeting discrete miRNA motifs provide a feasible, mutation-agnostic, and potentially generalizable therapeutic approach for haploinsufficient disorders. Collectively, these results advance both the mechanistic understanding of PKD1 regulation and the broader concept of targeted transcript derepression as a therapeutic modality.

## METHODS

### Generation of *Pkd1*^RC*17/RC^ mice

*Ksp*^Cre^;*Pkd1*^RC/RC^ female and male mice on a C57BL/6J background were used for creation of *Pkd1*^RC*17/RC^ mice^13, 23^. A DNA template was designed with 50-nucleotide homology arms flanking the 6 cardinal nucleotides complementary to the seed of miR-17, which were modified purine for purine and pyrimidine for pyrimidine as noted in Supplemental Table 4. A standard hormonal regimen was used to induce superovulation was induced in prepubertal female. Sperm was harvested from the epididymis of male mice, after which in vitro fertilization was performed. CRISPR reagents (HDR template, sgRNA and Cas9 protein, IDT) were introduced into the cytoplasm of one-cell fertilized eggs via electroporation using the Nepa21 Super Electroporator (NEPAGENE, Ichikawa, Japan). Eggs that survived electroporation were washed and cultured in microdrops of fresh M16 media. Next, the eggs were surgically delivered into oviducts of day 1 pseudopregnant ICR female mice. At postnatal day 27, identification of *Pkd1*^RC*17/RC^ mice was performed using genotyping and confirmed with Sanger sequencing. Primer sequences are noted in Supplemental Table 4.

### Mice

*Ksp*^Cre^, *Pkd1*^F/F^, and *Pkd1*^RC/RC^ mice on a C57BL/6J background were used for all experiments^13, 23, 24^. At the time of sacrifice, mice were anesthetized with isoflurane, blood was collected using cardiac puncture, and the right kidney was retrieved, weighed, and immediately flash-frozen in liquid nitrogen. The left kidney was perfused with 1x PBS followed by 4% (wt/vol) paraformaldehyde and then immersed in 4% paraformaldehyde for 2 hours before transferring to 1x PBS for storage or further processing. Equal male and female mice were used for all studies. All animal experiments have been approved by the UTSW Institutional Animal Care and Use Committee.

### Histology

Histology was performed using standard protocols by the UTSW Histology Core. The tissues were embedded in paraffin and sliced into 5µm sections. Staining was performed with hematoxylin-eosin (H&E) and unstained slides were used for immunofluorescence staining. All stained sections were imaged with a slide scanner. Cyst Index (cyst area/total kidney area) was calculated using Image J software.

### Isolation of primary kidney tubule epithelial cells

Primary epithelial cells were isolated from 10- to 14-day-old *Pkd1*^RC*17/RC*17^ *Pkd1*^RC*17/RC^, and *Pkd1*^RC/RC^ mice kidneys. Briefly, each mouse was perfused first with DMEM and then with DMEM containing 0.5% collagenase. Kidneys were retrieved, washed, and then minced into 1 mm pieces before incubating in prewarmed 0.5% collagenase in DMEM for 15 minutes at 37°C with shaking. Samples were passed through an 18-gauge needle (3-4x) and then returned to 37°C with shaking for an additional 15 minutes. Samples were subsequently filtered through a 40-µm cell strainer, washed with complete media, and pelleted. Cells were resuspended in PBS with 0.1% BSA then incubated in 10 µg/mL biotinylated DBA (VectorLabs #B-1035). After one hour, cells were pelleted and washed with PBS with 0.1% BSA, and subsequently DBA positive cells were isolated using CELLection Biotin binder kit (Invitrogen #11533D). Primary cells were plated and grown in culture media for 48 hours and then used for experiments or set up for immortalization. Primary *Pkd1*^RC/−^ and *Pkd1*^RC*17/−^ cells were immortalized using SV40 T Antigen Cell Immortalization Kit (ALSTEM #CILV01). Cells were placed in a 96-well plate for single clone expansion and genotyped for SV40, *Pkd1*^RC/−^ or *Pkd1*^RC*17/−^ and *Ksp*^Cre^ by DNA PCR. Clones positively identified by genotyping for SV40 and *Ksp*^Cre^ were stained for kidney tubule epithelial markers. Cell lines which stained positive for Aquaporin 2 (Sigma-Aldrich, #A7310) and HNF1β (Santa Cruz Biotechnology, #sc-22840) (Supplemental Figure 2) were used for subsequent analysis.

### qRT-PCR

Qiagen miRNeasy mini kit was used to extract total RNA. 0.5-1 µg of RNA was treated with DNase I (Invitrogen) and cDNA was produced with Invitrogen First Strand Superscript III cDNA synthesis kit. Q-PCR was performed using iQ SYBR Green Supermix (Bio-Rad). Each sample was loaded in duplicate or triplicate for analysis using the CFX ConnectTM Real-time PCR machine. 18s rRNA was used to normalize mRNA expression. For mature miRNA analysis, the Taqman MicroRNA Reverse Transcription Kit and Taqman Fast Advanced qPCR Mastermix (Thermo Fisher Scientific) were used, and values were normalized to U6. Primer sequences are noted in Supplemental Table 4. Primers for primary miR-17 were purchased from Thermo Fisher Scientific (Taqman Pri-miRNA-Assay #4427012).

### mRNA Stability Assay

Primary *Pkd1*^RC*17/RC^ cells were seeded into 6-well plates to reach 50 percent confluence. The next morning, culture media was replaced with media containing 5 µg/ml media of actinomycin. Cells were harvested at 0, 2, 4, 6 and 8 hours after actinomycin treatment for analysis. Similarly, normal collecting duct cells or *Pkd1*^RC/−^ cells were plated into 6 well plates (1.5 x 10^5^ cells per well) and the next morning transfected with control or *Pkd1* oligo using Lipofectamine 3000 (Invitrogen) to reach a final concentration of 40 nM. 48 hours after transfection, culture media was replaced with media containing 5 µg/ml media of actinomycin. Cells were harvested at 0, 2, 4, 6 and 8 hours after actinomycin treatment for analysis.

### Western blot

Total protein was extracted from kidneys by homogenization in a lysis buffer containing T-PER and protease phosphatase inhibitor (Thermo Fisher Scientific, #PIA32961). For in-vitro studies, adherent cells were scraped with cell scrapers in 1x PBS, transferred into 1.5 ml microcentrifuge tubes and centrifuged at 10000 rpm for 5 minutes. Supernatant was aspirated and then pellet was resuspended in T-PER+PPI. Protein concentrations were measured with Bradford Assay reagent. For proteins <150kDa in size, 10 µg was loaded for western blot. For full-length PC1 protein (462 kDa), 40-60 µg was used. For detection of PC1, protein lysate was prepared in 0.1M dithiothreitol and incubated at 25^°^C for 15 minutes immediately before gel electrophoresis. Samples were loaded onto NuPAGE 3-8% Tris-Acetate Protein Gel (Thermo Fisher Scientific) alongside a high molecular weight protein ladder (HiMark, Invitrogen) and run at 150V for 20 minutes at room temperature, followed by 40 minutes at 4^°^C. Protein transfer onto a nitrocellulose membrane was performed using the Invitrogen transfer system at 200 mAmps for 100 minutes at 4°C. The membrane was then blocked in 5% fat-free milk in TBST for 30 minutes and incubated in PC1 primary antibody (1:500 dilution) in TBST overnight. The next morning, the PC1 membrane was incubated on shaker for 1 additional hour at room temperature with PC1 primary antibody (1:250 dilution). Membranes were then washed thrice with TBST and incubated in secondary antibody (anti-mouse HRP-conjugated IgG at 1:500 for PC1) for 1 hour at room temperature. After 3 additional washes in TBST, membrane was developed using SuperSignal West Femto or Atto (Thermo Fisher Scientific) using Bio-Rad digital imager. For actin for PC1 blots, 3 µl of the same protein lysate prepared in 0.1M dithiothreitol was loaded onto mini-PROTEAN SDS-polyacrylamide precast gels (Bio-Rad), alongside a standard molecular weight ladder (Precision Plus Protein Kaleidoscope Standards, Bio-Rad). Protein transfer and all subsequent steps were identical as for all other western blots described below.

For all other western blots, protein was prepared 4X NuPage LDS Sample buffer with 0.5% β-mercaptoethanol. For OXPHOS assessment, samples were kept at 50°C for 10 minutes before loading. For all other proteins, samples were boiled for 3 minutes at 98°C and cooled to room temperature, before loading onto mini-PROTEAN SDS-polyacrylamide precast gels (Bio-Rad), against a standard molecular weight ladder (Precision Plus Protein Kaleidoscope Standards, Bio-Rad). Gels were run at 200V at room temperature until the dye ran out. Protein transfer was performed using the Trans-Blot Turbo Transfer System (Bio-Rad) with the mixed MW or Mini TGX program. Nitrocellulose membranes were then blocked with 5% fat-free milk in TBST for an hour, and then probed with primary antibody (1:1000) for 1 hour at room temperature followed by overnight at 4°C. After completing incubation with primary antibody, membranes were washed three times with TBST, and then probed for 1 hour at room temperature with secondary antibody (1:5000). For β-actin, an HRP-conjugated actin antibody diluted at 1:40000 (Sigma Aldrich) was used, with incubation of 1 hour. Blots were then washed three times and developed using the chemiluminescence substrate Pierce ECL, SuperSignal West Dura or Femto (if needed) and imaged with the Bio-Rad digital imager. Quantification was performed using ImageJ.

Primary antibodies used: PC1 (7E12 Santa Cruz, #sc-130554), PC2 (PKD-RRC), AQP2 (Sigma-Aldrich, #A7310), Beta-Actin (Sigma Aldrich, #A3854), Mannose Receptor (Abcam, #ab64693), Alpha Smooth Muscle (Sigma Aldrich, #F377), Phospho-mTOR (Cell Signaling, #2971), pCREB (Cell Signaling, #9198), Total OXPHOS Rodent WB Antibody Cocktail (Abcam, #ab110413), c-Myc (Abcam, #ab185656). Secondary Antibodies used: goat-anti-rabbit IgG HRP-conjugated and goat-anti-mouse IgG HRP-conjugated (Thermo Fisher Scientific).

### Immunofluorescence

Paraffin-embedded kidney tissue slides were deparaffinized by incubating at 60°C for 1 hour, followed 3 serial 5-minute washes in Histo-clear (Thermo Fisher Scientific). Slides were rehydrated by washing in 100%, 95%, and 70% ethanol for 5 minutes each followed by direct immersion in 1x PBS. Antigen retrieval was performed using warm sodium citrate. After cooling, slides were blocked in blocking solution (1x PBS + 10% goat serum + 0.1% BSA) for 1 hour at room temperature and then incubated in primary antibody at 1:500 dilution overnight. The next morning, slides were washed thrice with 1X PBS and then incubated in secondary antibody at 1:500 dilution for 1 hour at room temperature in a light-protected slide box. Slides were again washed with 1X PBS three times and then mounted with Vecta Shield containing Dapi. Zeiss AxioObserver deconvolution microscope and Zeiss Axioscan Z1 microscope were used for imaging. Primary antibodies used: phospho-histone H3 (Sigma Aldrich, #H0412). Secondary antibodies: Alexa Fluor 594 Dye (Fisher Scientific, #PIA32758).

### Proteomics

Protein was prepared in 0.1M dithiothreitol, incubated at 25°C for 15 minutes, then immediately loaded into mini-PROTEAN TGX 4-10% SDS-polyacrylamide gels (Bio-Rad). Gels were run at 180V for 5-10 minutes to allow protein to enter the top 1 cm of the gel. The gel was then removed from its cassette and placed in Coomassie Blue R-250 (Bio-Rad) on a shaker for 90 minutes. The stained gel was washed in ddH_2_O and the top 1 mm of each well was discarded to reduce variability due to captured detergents. The next 1 cm of each well was excised and diced into 1 mm cubes for protein extraction prior to LC-MS. Gel protein extraction, trypsin digestion, and complex mixture LC-MS using a 90-minute HPLC gradient were performed by the Proteomics Core at UTSW to identify proteins. Abundance of proteins was normalized to beta-actin for each sample. Three samples were analyzed for each genotype. One-way Anova was used to identify differentially abundant proteins. Differential pathway analysis was performed using Ingenuity Pathway Analysis software.

### Magnetic Resonance Imaging

84-day old mice were imaged with MRI at the UTSW Pre-Clinical MRI Research Core on the 7-T Bruker Biospec scanner. Kidney volumes were quantified manually by multiplying areas of kidneys for each slice by the slice thickness, using PiPro provided by the UTSW Pre-Clinical MRI Research Core.

### Serum creatinine

Serum creatinine was measured using capillary electrophoresis by the UTSW Division of Nephrology.

### Oligonucleotide Design

Binding regions for oligonucleotide candidates were found using the findMotif utility by Kent Informatics, distributed by UCSC. This search was conducted against the T2T-CHM13 v2.0 genome for human, and GRCm39 genome for mouse. Gene annotations were added by intersecting the binding motifs with NCBI Refseq annotations using bedtools. Monomer and homodimer free energies were calculated using the RNAcofold tool from the ViennaRNA package. Optimal LNA placement and oligonucleotide synthesis was performed by Qiagen.

### Luciferase Assay

mIMCD3 or *Pkd1*^RC/−^ cells were seeded into six-well dishes (2 × 10^5^ cells per well) and transfected with 0.4 μg of pLS-*Pkd1*-3′-UTR plasmid, 10 nM of miR-17 or scramble mimic (Dharmacon) and 40 nM of Control or *Pkd1* oligo (Qiagen). Cells were also transfected with 0.04 μg of the pGL3-Control plasmid (Promega Corp) encoding Photinus luciferase to serve as control for differences in transfection efficiency. Lipofectamine 2000 (Invitrogen) was used as a transfection reagent. After forty-eight hours, the cells were lysed in 250 μl of passive lysis buffer (Promega Corp), and 40 μl of the cell lysate was added to 96-well plates. Photinus and Renilla luciferase activities were measured by using the Dual-Luciferase Reporter Assay System (Promega Corp) according to the manufacturer’s directions.

### Cell Lines

*Pkd1*^RC/−^ cells, kidney epithelial cells, and mIMCD3 cells were used in these experiments. mIMCD3 cells were obtained from ATCC. Collecting duct derived kidney epithelial cells and *Pkd1*^RC/−^ cells are immortalized tubule derived kidney epithelial cells derived in our laboratory from 12-day old male mouse kidneys^5, 25^.

### Human ADPKD Cell *Lines*

WT9-7 cells (*PKD1*.Q2556X) were obtained from ATCC. Immortalized human ADPKD cell lines were derived from primary human cyst cells from human kidneys affected by ADPKD obtained from the PKD Research Biomarker and Biomaterial Core at University of Kansas Medical Center. The use of tissues was approved by the KUMC Institutional Review Board and complied with federal regulations. Mutation analysis has been previously performed by Ambry Genetics (Aliso Viejo, CA)^5^. Culture media consisted of DMEM/F12 + + (Gibco, #10565–018) supplemented with 10% FBS, 5 μg/kg insulin, 5 μg/mL transferrin, and 5 ng/mL sodium selenite. Each primary cell line was grown in an atmosphere of 95% air and 5% CO_2_ at 37 °C until 80 percent confluency. At the second passage primary cell lines underwent immortalization as described above. Single clones which expressed SV40 were selected for expansion and downstream utilization. Immortalized human ADPKD cell lines were grown to 80% confluency, then cells were trypsinized and seeded at 2 x 10^5^ cells per well in a 6-well plate. The next day, cells were transfected with Lipofectamine 3000 + control oligo, P1 oligo or P2 oligo. After 48 hours, cells were harvested or fixed with 100% methanol for analysis or placed in Matrigel for cystogenesis assay.

### 3D Cystogenesis Assay

96-well plates or 8-well chamber slides and 200 µl sterile pipette tips were pre-cooled at −20°C for a minimum of 6 hours. The floor of each well of an 8-well chamber slide or selected wells of a 96-well plate was carefully coated with 25 µl of 100% Matrigel (Fisher Scientific, #354234) using 200 µl sterile pipette tips. The plate was then placed in a 37°C incubator for 30 minutes to set the Matrigel. During this time, cells were washed with 1X PBS, trypsinized and filtered through a 40 µm cell strainer to create a single-cell suspension. Cells were counted with Hemocytometer and diluted to reach a final concentration of 5000 cells per 150 µl. 150 µl of cell suspension was combined 1:1 with 4% Matrigel and placed in each well. Each treatment condition was seeded in triplicate and incubated at 37°C for 7 days. After 72 hours, each well was supplemented with 100 µl of epithelial media. On day 7, the cysts were imaged using a Leica DMI 3000B light microscope. The images were analyzed using ImageJ software to measure cyst size. In immortalized cells, each assay was repeated three times.

### Immunofluorescence Staining of Cells

Adherent cells were fixed onto chamber slides with ice-cold 100% methanol for 5 minutes at 4°C and then washed thrice with 1X phosphate buffered saline. The cells were then blocked for 30 minutes to 1 hour in 1X PBS + 10% goat serum + 0.1% BSA + 0.1 M glycine + 0.1% Tween 20 (blocking solution) at room temperature. pCREB antibody (Cell Signaling #9198) at 1:400 dilution in blocking solution was added and slides were incubated overnight at 4°C. The next morning, primary antibody was removed and slides were washed 3x in 1X PBS. Then secondary antibody was applied at 1:400 dilution for 1 hour at room temperature and then again washed thrice in 1X PBS. Cells were counterstained with DAPI diluted 1:10,000 in 1X PBS and visualized using Carl Zeiss AxioObserver microscope. All conditions for each experiment were processed and imaged simultaneously.

### MitoTracker staining

Cells were washed with 1X PBS and then incubated in 100 nM MitoTracker Red CMXRos dissolved in serum free DMEM (Thermo Fisher Scientific). After eight minutes the media was aspirated and replaced with regular epithelial media and cells were immediately imaged using Carl Zeiss AxioObserver microscope. All images for each experiment were taken with identical exposure to allow comparison of intensity of MitoTracker fluorescence, a proportional surrogate of mitochondrial membrane potential.

### RNA Sensor assay

CSM vector pDAC565 (Addgene #195242) was obtained from Addgene with institutional MTA. We followed the creators’ instructions to design and insert a crRNA specific to *Pkd1* (Supplementary Table 4) into the CSM vector^14^. To image *Pkd1* mRNA in live cells, 2×10^4^ cells were seeded into 24 well plates. The next morning, cells were transfected with 0.4 µg of CSM plasmid and control oligo or *Pkd1* oligo attotal concentration of 40 nM using Lipofectamine 3000. At 48 hours, cells were evaluated using the Zeiss AxioObserver microscope at 20x magnification. 200 cells per condition which exhibited green background signal (indicating plasmid transfection) were evaluated for the presence of GFP fluorescent puncta, which marked *PKD1* mRNA. The percentages of cells positive for *Pkd1* mRNA puncta were recorded. Each experiment was repeated at least three times.

### Statistical Analysis

Equal males and females were used for all studies. Two-tailed Students *t-test* was used to compare 2 groups. Analysis of variance (ANOVA) followed by Tukey’s multiple comparison test was used to compare 3 or more groups. Surivival analysis was determined by Mantel Cox test. All statistical analysis was performed using GraphPad Prism software, Python scipy modulator Microsoft Excel. P<0.05 was considered statistically significant.

## Data Availability

Proteomics dataset is available at MassIVE: Accession number: MSV000099490.

## Acknowledgements

This work is supported by the National Institute of Health (5R01DK102572 and 5R01DK133186) to VP and (1R01DK139033) to RL. This work is also supported by PKD-RRC Sprint Challenge Award to RL.

## Contributions

RL, CS, LB, MZ, JA, AS, PC, HR, and VP designed and executed experiments and interpreted results. RL, CS, LB, and VP designed the figures. RL, CS, and VP wrote the manuscript with constructive input from all authors.

## Competing Interests

V.P. has served as a scientific consultant for Otsuka Pharmaceuticals, Maze Therapeutics, Travere Therapeutics, and Regulus Therapeutics. V.P. serves as the chair of the Scientific Advisory Panel for the PKD Foundation. V.P. lab has a sponsored research agreement with Regulus Therapeutics, a Novartis company, which is unrelated to this work. V.P. has licensed patents (US11168325B2) involving anti-miR-17 for the treatment of ADPKD, which is unrelated to the current manuscript. A patent application (PCT/US2024/017086) has been filed by UT Southwestern Medical Center (RL, LB, MZ, HR, and VP listed as authors) which includes some of the findings presented in this manuscript.

---

**Supplementary Figure 1.**
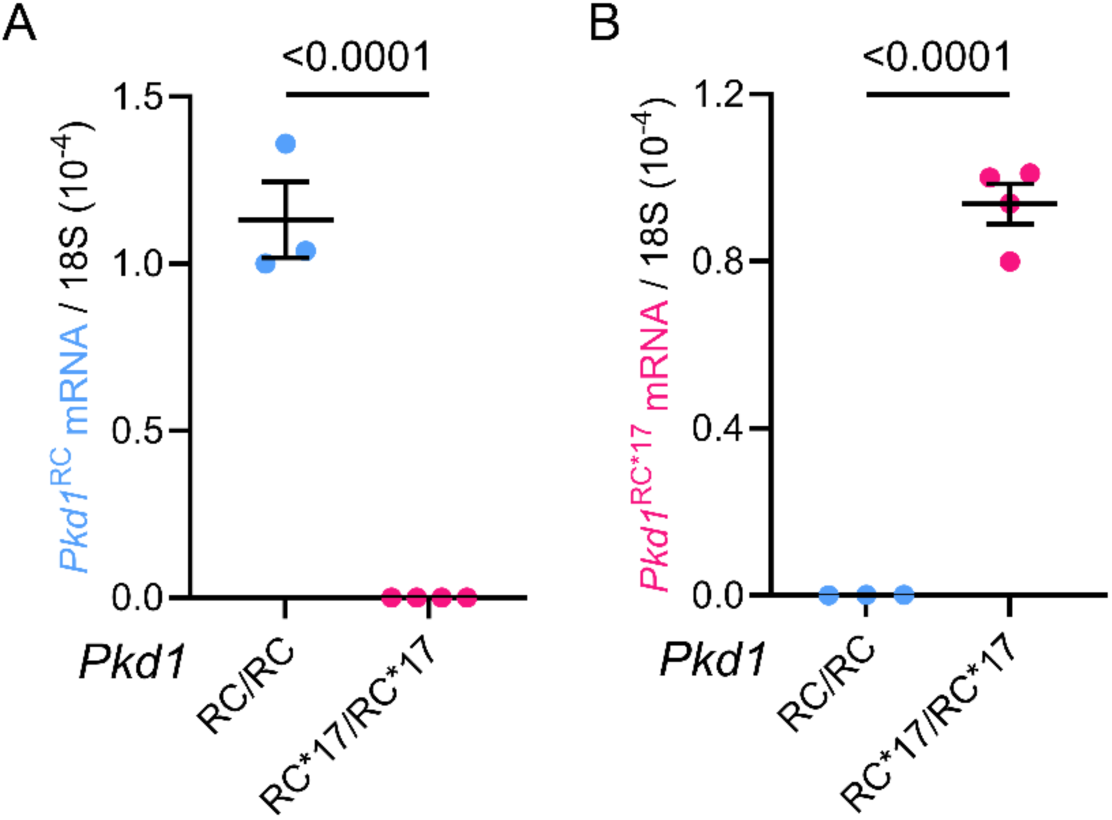
Validation of allele-specific qRT-PCR primers to detect *Pkd1*^RC^ and *Pkd1*^RC*17^ alleles. **A&B.** qRT-PCR using RNA from primary kidney epithelial cells derived from *Pkd1*^RC/RC^ and *Pkd1*^RC*17/RC*17^ mice demonstrates allele-specific amplification of *Pkd1*^RC^ and *Pkd1*^RC*17^ transcripts respectively. Error bars indicate SEM. N=3, biological replicates. Statistics: Unpaired, two-tailed, t-test.

**Supplementary Figure 2.**
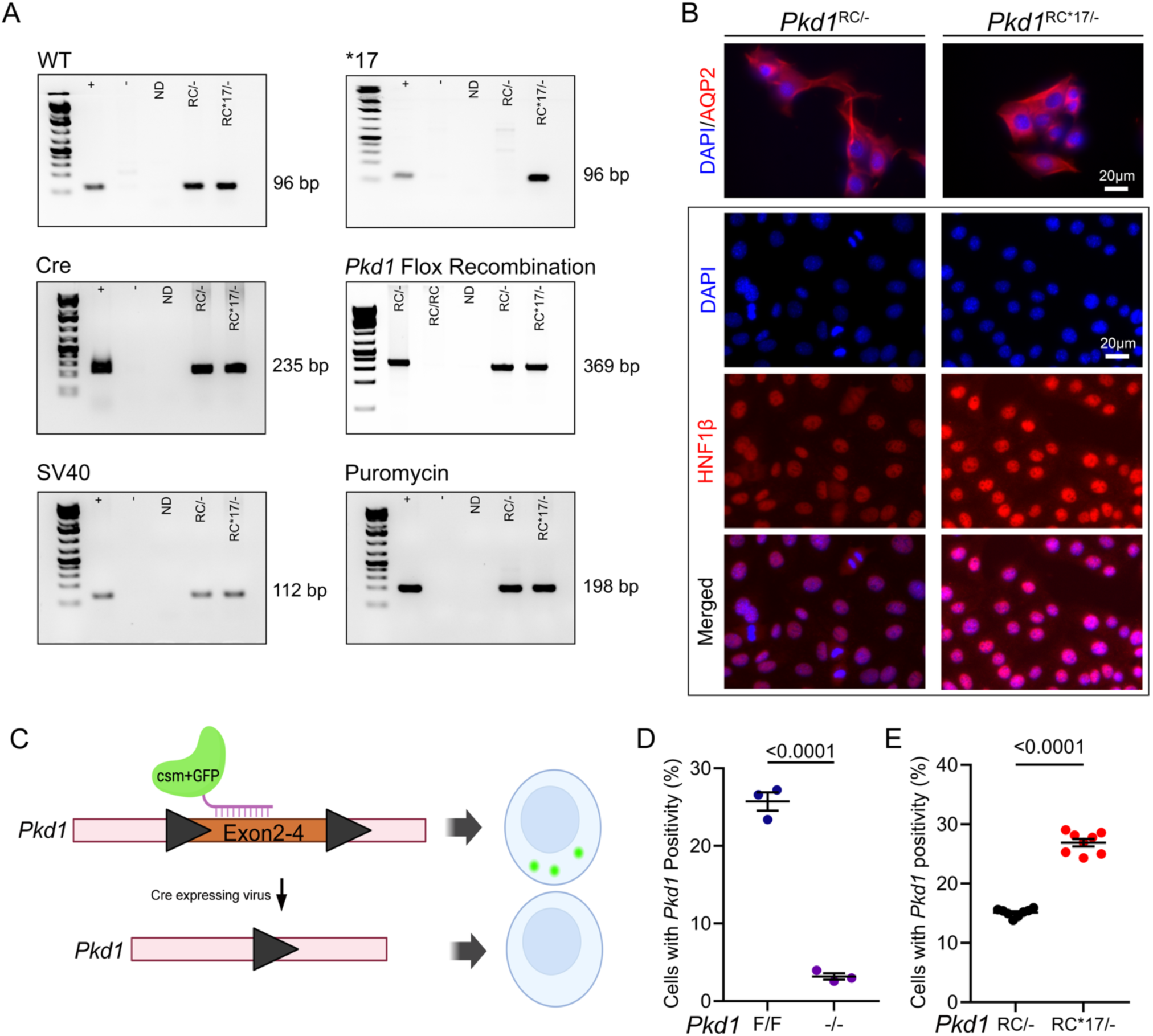
Characterization of *Pkd1*^RC/−^ and *Pkd1*^RC*17/−^ cell lines. Kidney epithelial cells enriched for DBA were isolated from 10-day-old *Ksp*^Cre+^;*Pkd1*^RC/F^ and *Ksp*^Cre+^;*Pkd1*^RC*17/F^ mice kidneys. Single clones were isolated for subsequent analysis. **A.** Gel electrophoresis confirmed genotypes of *Pkd1*^RC^ (WT 3’UTR) and *Pkd1*^RC*17^ (*17 3’UTR) alleles in *Pkd1*^RC/−^ and *Pkd1*^RC*17/−^ cell lines. Presence of *Ksp*Cre transgene and successful recombination of *Pkd1* is shown. Immortalization with SV40 is confirmed by the presence of SV40 and puromycin cassette. **B.** Immunofluorescence demonstrated presence of collecting duct marker aquaporin 2 (AQP2) and epithelial marker HNF1B. **C-D.** Schematic illustration of the CRISPR-based live-cell RNA sensing system to detect *Pkd1* mRNA. A sgRNA was designed specifically to target *Pkd1* exons 2-4. In *Pkd1*^F/F^ cells, the *Pkd1*-SgRNA directs the catalytically inactive CRISPR-Cas complex (csm-GFP) to the mRNA, resulting in fluorescent puncta. However, *Pkd1*^−/−^ cells (derived from *Pkd1*^F/F^ cells by infecting them with a virus expressing Cre recombinase) should demonstrate no fluorescent puncta signal with the csm-GFP-*Pkd1*-SgRNA system. Accordingly, live cell quantification demonstrates GFP puncta in *Pkd1*^F/F^ cells but not *Pkd1*^−/−^ cells, implying specificity of *Pkd1*-SgRNA in detecting *Pkd1* mRNA. **E.** This *Pkd1* mRNA detection system was used in *Pkd1*^RC*17/−^ and *Pkd1*^RC/−^ cells (shown in A-B). *Pkd1* mRNA detection was 50% higher in in *Pkd1*^RC*17/−^ compared to *Pkd1*^RC/−^ cells. Error bars indicate SEM. N=3, biological replicates. Statistics: Unpaired, two-tailed, t-test.

**Supplementary Figure 3.**
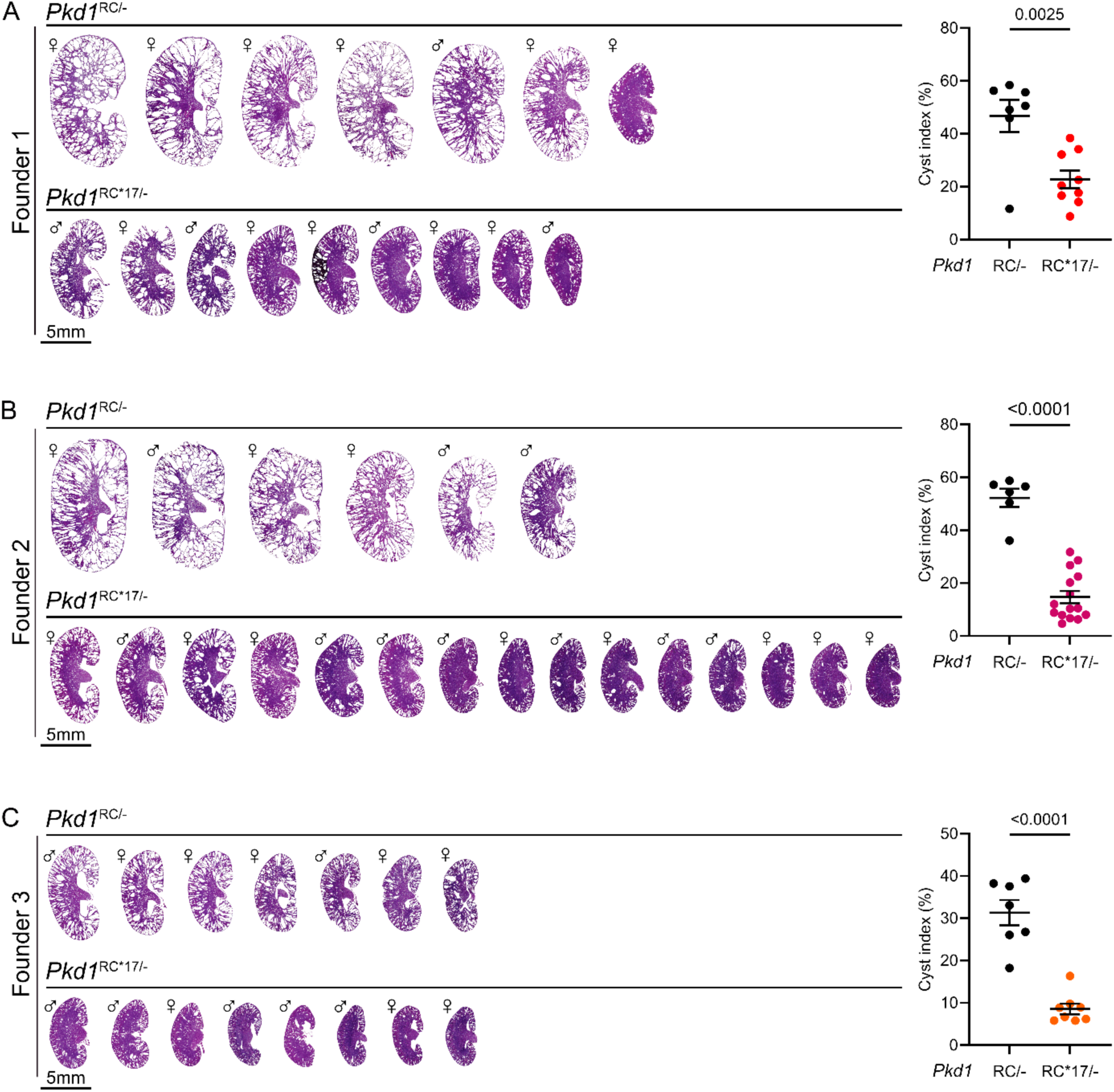
Kidney section H&E and cyst indices of *Pkd1*^RC/−^ and *Pkd1*^RC*17/−^ mice. Gross H&E of kidney sections and the associated kidney cyst index of every 18-day-old *Pkd1*^RC/−^ and *Pkd1*^RC*17/−^ mouse analyzed in this study is shown. The data from Founder 1, 2, and 3 are shown in A, B, and C, respectively. Gender is indicated by symbol to the left of each kidney image. Error bars indicate SEM. Statistics: Unpaired, two-tailed, t-test.

**Supplementary Figure 4.**
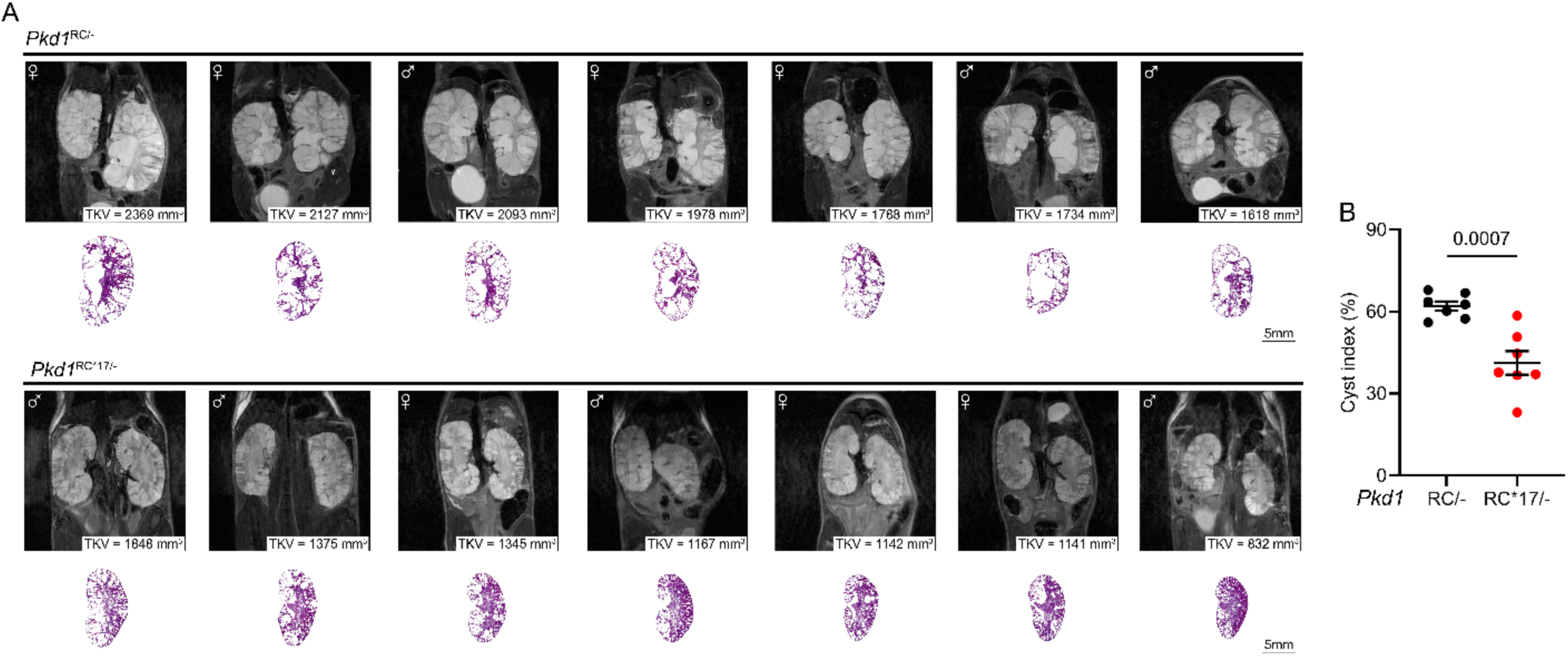
MRI-based anatomical and kidney histological analysis of aged *Pkd1*^RC/−^ and *Pkd1*^RC*17/−^ mice. **A.** MRI of midline coronal section of kidneys with corresponding left kidney section H&E of every 84-day-old littermate *Pkd1*^RC/−^ and *Pkd1*^RC*17/−^ mice analyzed in this study is shown. Gender of each mouse is indicated in top left of each MRI image. Corresponding total kidney volume (TKV) is denoted in mm^3^ in each MRI. **B.** Cyst index based on kidney section H&E in *Pkd1*^RC*17/−^ kidneys compared to *Pkd1*^RC/−^ kidneys is shown. Error bars indicate SEM. Statistics: Unpaired, two-tailed, t-test.

**Supplementary Figure 5.**
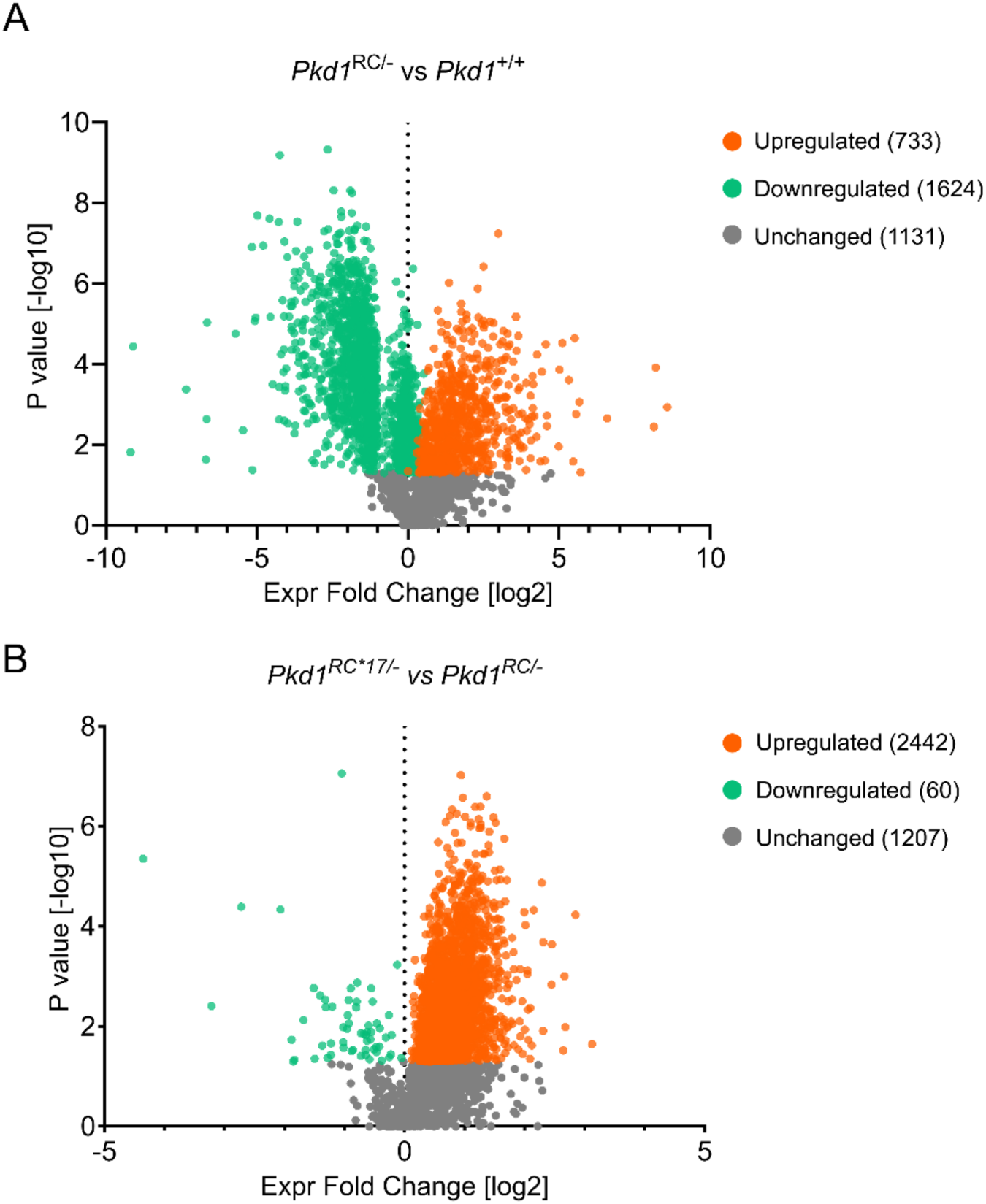
Proteomic analysis of *Pkd1*^+/+^, *Pkd1*^RC/−^, and *Pkd1*^RC*17/−^ kidneys. **A&B**. Volcano plot depicts change in protein expression patterns (upregulated = orange; downregulated = green; unchanged = gray) between *Pkd1*^+/+^ and *Pkd1*^RC/−^, and *Pkd1*^RC^ and *Pkd1*^RC*17/−^ mice kidneys. P < 0.05 was considered significantly changed. N=3 biological replicate for each group. Statistics: Unpaired, two-tailed, t-test.

**Supplementary Figure 6.**
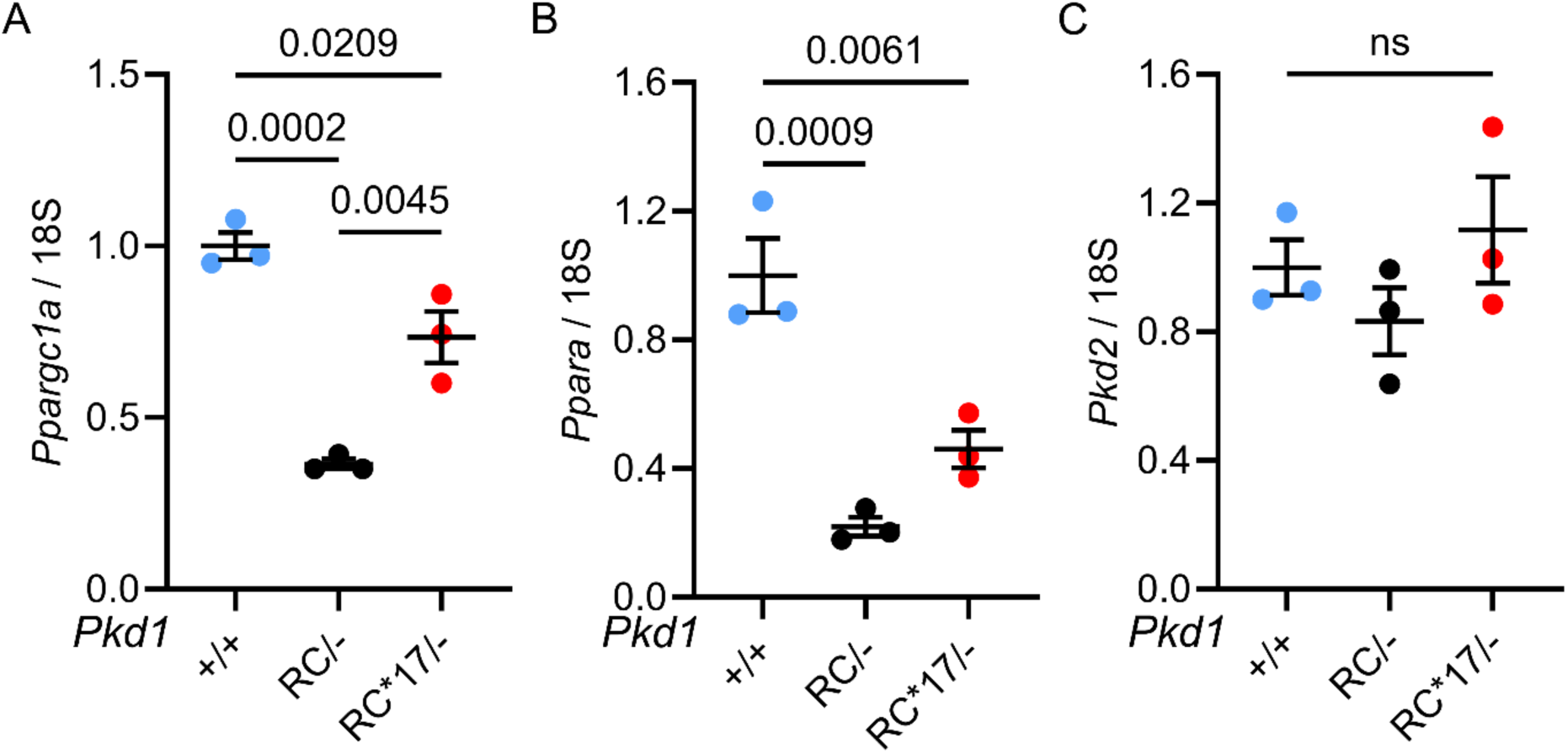
qRT-PCR of additional genes involved in the c-Myc-miR-17 axis. **A.** Expression of *Ppargc1a*, a key regulator of mitochondrial biogenesis is reduced in *Pkd1*^RC/−^ kidneys compared to *Pkd1*^+/+^ kidney, but is restored in *Pkd1*^RC*17/−^ kidneys. **B.** In contrast, the expression of *Ppara*, a direct target of miR-17 in the context of PKD, is not improved in *Pkd1*^RC*17/−^ kidneys. **C.** The expression of *Pkd2*, another miR-17 target, was also not different between *Pkd1*^+/+^, *Pkd1*^RC/−^ and *Pkd1*^RC*17/−^ kidneys. Error bars indicate SEM. N=3 biological replicates in all groups. Statistics: ANOVA with Tukey’s.

**Supplementary Figure 7.**
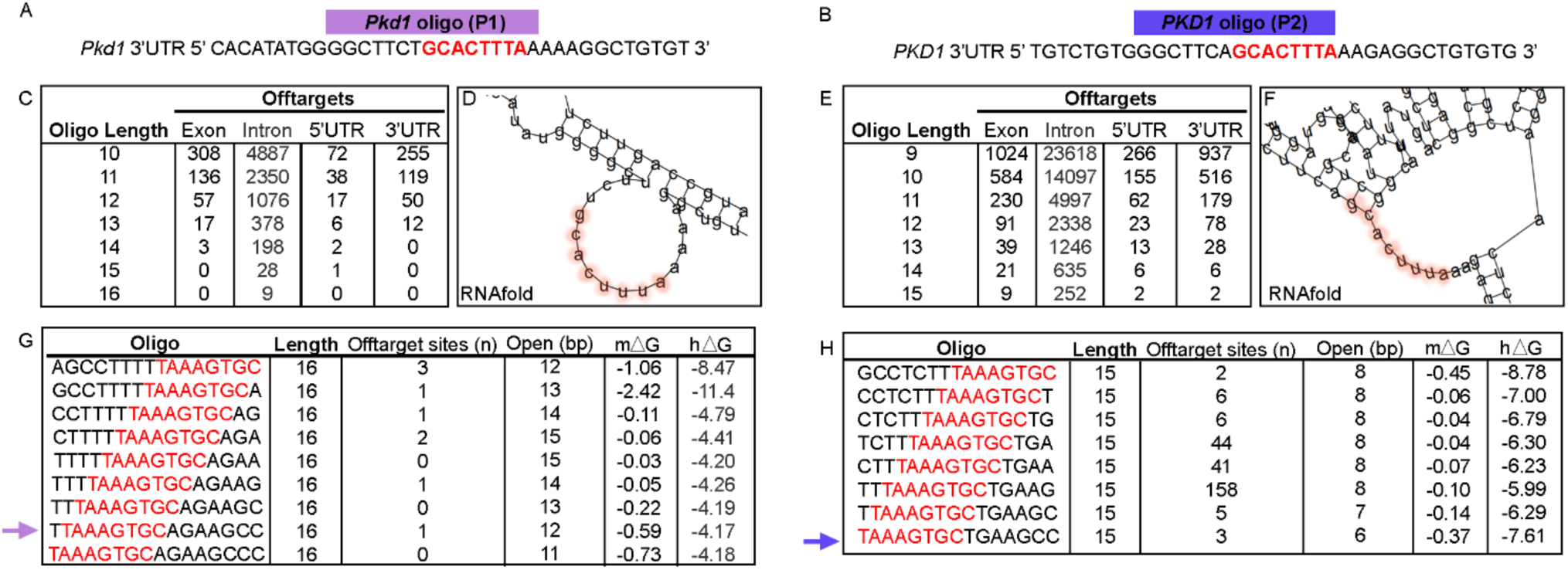
Consideration for design of miR-17 binding motif targeting ASOs. **A&B.** Genomic sequence of mouse and human *PKD1* 3’UTR is shown with nucleotides highlighted in red indicating the miR-17 binding motif. **A.** The exact region for the mouse-specific *Pkd1* oligo (referred to as P1 oligo in the main text) binding is shown above in purple. **B**. The exact region for the human-specific *PKD1* oligo (referred to as P2 oligo in the main text) binding is shown above in blue. **C&E**. Bioinformatic analysis of proposed oligo lengths and potential off-target binding is noted. The shorter oligonucleotide lengths were predicted to have numerous off-target binding. This analysis for the P1 oligo is shown in C, and P2 is shown in E. **D&F.** RNA fold was used to examine the predicted secondary structure of the *PKD1* 3’-UTR. The nucleotides highlighted in pink comprise the miR-17 binding motif. **G&H.** After determination of optimal length, a tiling approach was used to identify the ASO with the least number of off targets and minimal potential for hair-pinning (monomer delta G, mΔG) or self-dimerization (homodimer delta G, hΔG). The oligos marked with arrows were chosen for further experimentation.

**Supplemental Figure 8.**
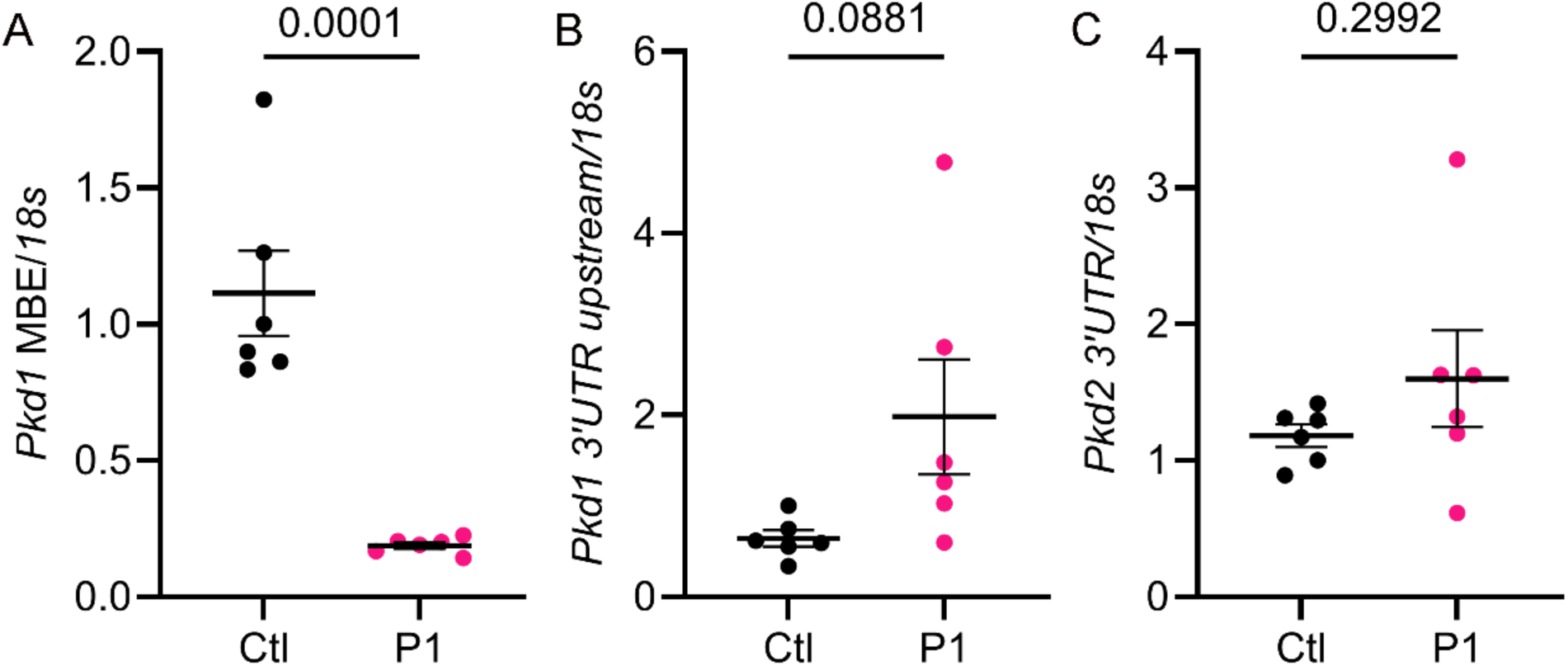
P1 oligo specifically binds to the miR-17 motif in *Pkd1* 3’UTR. Murine kidney epithelial cells were transfected with 40 nM P1 oligo or control oligo (Ctl). After 48 hours, cells were harvested and RNA was extracted for target engagement analysis. **A.** qRT-PCR demonstrates no amplification of *Pkd1* mRNA when probed at the *Pkd1*-miR17 motif in cells treated with P1 compared to Ctl oligo, indicating interference due to P1 oligo binding. **B**. However, *Pkd1* mRNA is detected upstream of the oligo binding site, implying specificity of P1 binding to the 3’-UTR motif. **C.** *Pkd2* is direct miR-17 target and contains a conserved miR-17 motif in its 3’UTR. No change in abundance of *Pkd2* mRNA was observed when probed directly at its miR-17 binding site, indicating specific binding of P1 to *Pkd1* 3-UTR but not to the closely related *Pkd2* 3’UTR. Error bars indicate SEM. N=6 biological replicates in all group. Statistics: Unpaired, two-tailed, t-test.

**Supplemental Figure 9.**
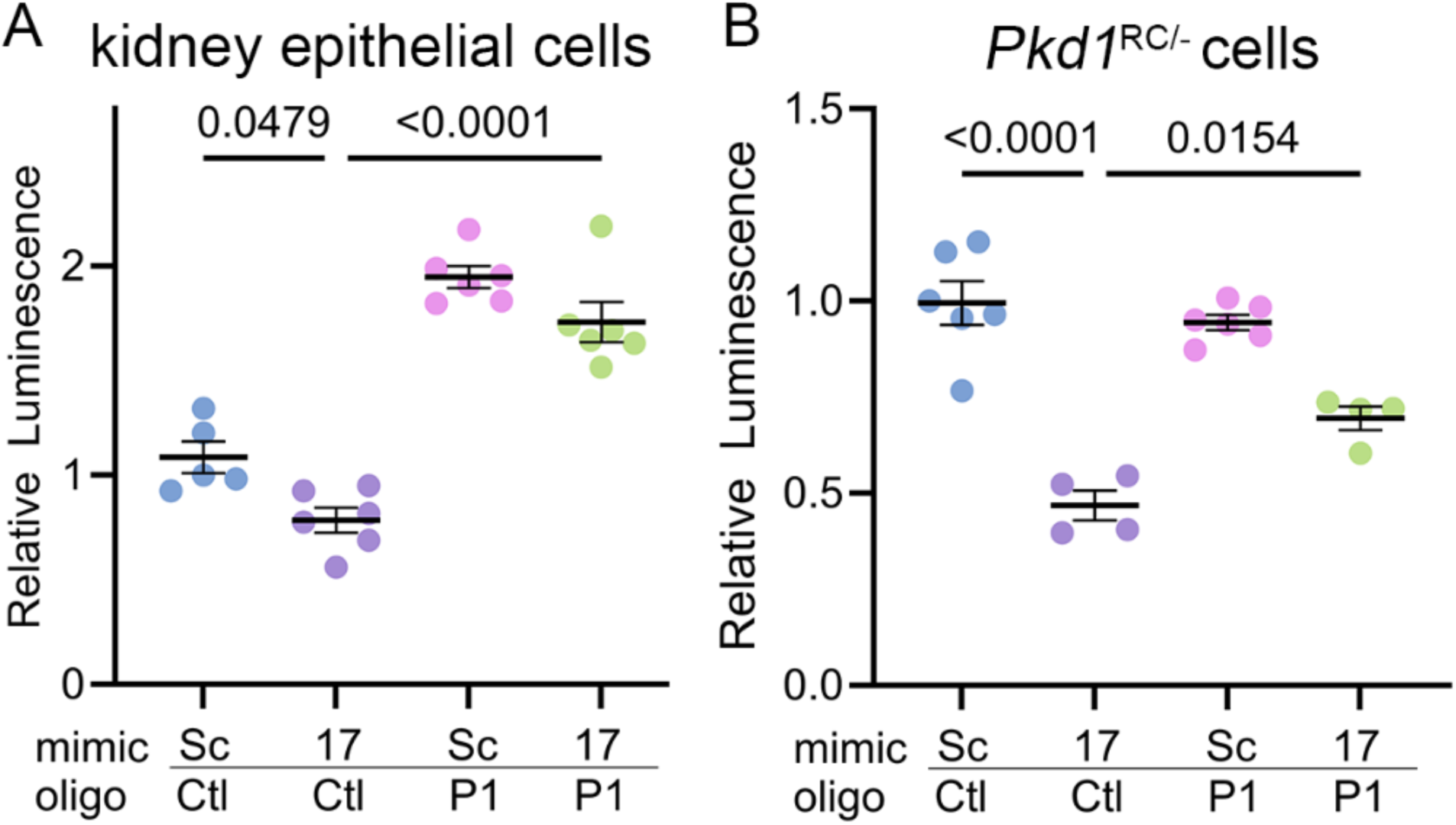
Functional engagement of the *Pkd1* 3’UTR by P1 oligo. **A&B**. Murine kidney epithelial cell lines were transfected with *Pkd1* 3’UTR luciferase reporter plasmid, miR-17 or scramble mimic, and P1 or control oligo. Luciferase reporter assay demonstrates repression of luminescence in the setting of miR-17 mimic which is recovered in the presence of P1 oligo. Error bars indicate SEM. Each data point is biological replicate with an average of 3 technical replicates. Statistics: ANOVA with Tukey’s.

**Supplemental Figure 10.**
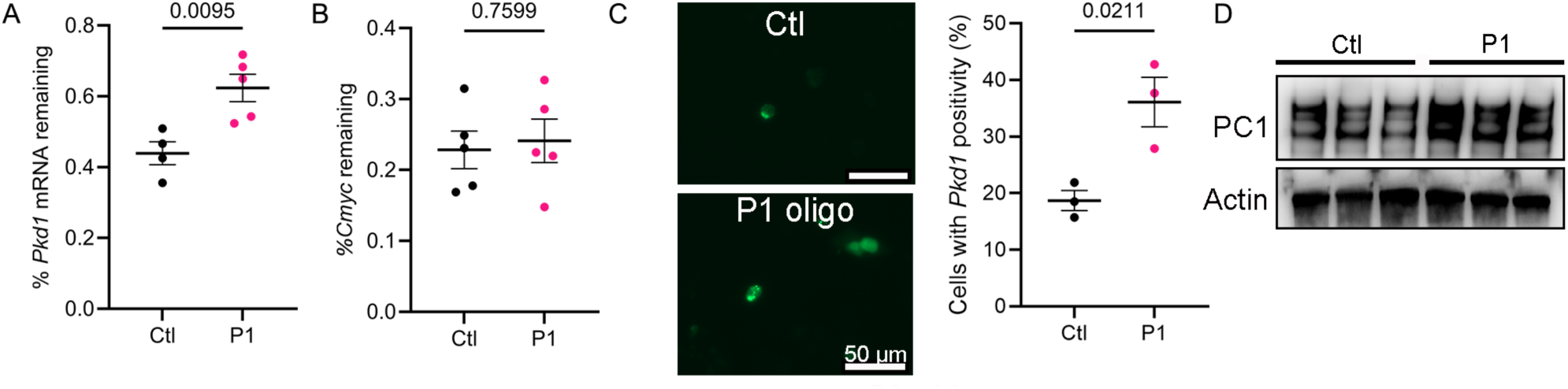
P1 oligo stabilizes endogenous *Pkd1* mRNA and increases PC1 protein. **A-B**. Collecting duct-derived kidney epithelial cells treated with control (ctl) oligo or P1 oligo for 48 hours and were subsequently treated with Actinomycin D for 6 hours to arrest de-novo transcription. **A**. qRT-PCR demonstrates 30% more *Pkd1* mRNA in cells treated with P1 oligo compared to control oligo treated cells, implying improved *Pkd1* transcript stability in P1-treated cells. **B.** As a negative control, the abundance of c-Myc mRNA remains unchanged**. C.** Collecting duct-derived kidney epithelial cells treated with control (ctl) oligo or P1 oligo for 48 hours and co-transfected with csm-GFP-*Pkd*1-SgRNA system for live-cell detection of *Pkd1* mRNA transcript. Immunofluorescence images and subsequent quantification show that cells treated P1 oligo exhibit increased *Pkd1* mRNA signal compared to Ctl-oligo treated cells. **D.** Western blot shows PC1 protein is increased in P1-treated kidney epithelial cells compared to Ctl-oligo treated cells. Error bars indicate SEM. Statistics: Unpaired, two-tailed, t-test. Actin serves as the loading control.

